# A flexible loop in the paxillin LIM3 domain mediates direct binding to integrin β3

**DOI:** 10.1101/2023.01.26.525744

**Authors:** Timo Baade, Marcus Michaelis, Andreas Prestel, Christoph Paone, Nikolai Klishin, Laura Scheinost, Ruslan Nedielkov, Christof R. Hauck, Heiko M. Möller

**Author notes:** Address correspondence to: Heiko M. Möller, Universität Potsdam, Institut für Chemie, Karl- Liebknecht- Straße 24-25, D-14476 Potsdam, Christof R. Hauck, Lehrstuhl Zellbiologie, Fachbereich Biologie, Maildrop 621, Universität Konstanz, Universitätsstrasse 10, 78457 Konstanz, Germany. Section for Biomolecular Sciences, The Kaj Ulrik Linderstrøm-Lang Centre for Protein Science, Structural Biology and NMR laboratory, Ole Maaløes Vej 5, room 3-0-37, DK-2200 Copenhagen N. Authors contributed equally (Timo Baade, Marcus Michaelis). Email addresses: TB MM AP CP NK LS RN CRH HMM.

## Abstract

Integrins are fundamental for cell adhesion and the formation of focal adhesions (FA). Accordingly, these receptors guide embryonic development, tissue maintenance and haemostasis, but are also involved in cancer invasion and metastasis. A detailed understanding of the molecular interactions that drive integrin activation, focal adhesion assembly, and downstream signalling cascades is critical. Here, we reveal a direct association of paxillin, a marker protein of focal adhesion sites, with the cytoplasmic tails of the integrin β1 and β3 subunits. The binding interface resides in paxillin’s LIM3 domain, where based on the NMR structure and functional analyses a flexible, seven amino acid loop engages the unstructured part of the integrin cytoplasmic tail. Genetic manipulation of the involved residues in either paxillin or integrin β3 compromises cell adhesion and motility. This direct interaction between paxillin and the integrin cytoplasmic domain identifies an alternative, kindlin-independent mode of integrin outside-in signalling particularly important for integrin β3 function.

## Introduction

Integrins are specialized cell surface receptors of animal cells that sense the extracellular matrix and coordinate cell adhesion with the organization of the cytoskeleton (Sun et al., 2016). Integrins are transmembrane glycoproteins consisting of an α and β subunit, with 18 distinct integrin α and 8 integrin β subunits encoded in the human genome (Hynes, 2002). The high affinity, active conformation of these heterodimers can be stabilized either by extracellular ligand binding (outside-in signalling) or by distinct intracellular signalling processes, which allow the association of the cytosolic scaffolding proteins talin and kindlin with the β subunit cytoplasmic tail (inside-out signalling) (Sun et al., 2019). Active integrins together with their binding partners talin and kindlin serve as the nucleus for the initiation of large, multimeric protein complexes termed focal adhesions (FAs) (Geiger and Yamada, 2011; Moser et al., 2009). Combined biochemical, genetic and microscopic analyses have revealed the stratified layout of focal adhesions and identified the characteristic compendium of signalling and adaptor proteins, the so-called integrin adhesome (Chastney et al., 2020; Horton et al., 2015; Kanchanawong et al., 2010; Zaidel-Bar et al., 2007). While the essential roles of talin and kindlin in initiating integrin-based adhesion sites in various cell types have become clear (Bachmann et al., 2019), the function of other core adhesome proteins during the initial steps of focal adhesion formation is still debated.

For example, LIM (Lin-11, Isl1, MEC-3) domain containing adapter proteins are a highly enriched subgroup of integrin adhesome proteins thought to be involved in mechanosensing (Anderson et al., 2021; Horton et al., 2015; Schiller et al., 2011). Individual LIM domains encompass ∼60 amino acids forming a double zinc finger motif, which mediates binding to other proteins or nucleic acids (Kadrmas and Beckerle, 2004; Matthews et al., 2009). A prominent member of this group of adapter proteins is paxillin, which contains four LIM domains and is ubiquitously expressed in mammalian tissues (Deakin et al., 2012; Deakin and Turner, 2008). Paxillin is commonly employed as a marker for FAs and nascent focal complexes under various conditions, even where the normal morphology, function and architecture of FAs is disturbed. Paxillin is one of the first proteins recruited to FAs (Digman et al., 2008), efficiently localizes there even in the absence of myosin-generated forces (Pasapera et al., 2010), and can recruit vinculin to FAs in talin knockout cells (Atherton et al., 2020). Paxillin is found in nanometer distance from the plasma membrane with the carboxy-terminus detected in the same confined membrane-proximal layer as the cytoplasmic domain of integrin αv (Kanchanawong et al., 2010). A main determinant of paxillin’s efficient recruitment to integrins seems to be its association with kindlin2, which has been mapped to the amino-terminal LD domains and the carboxy-terminal LIM4 domain of paxillin (Bottcher et al., 2017; Gao et al., 2017; Zhu et al., 2019). Interestingly, initial work identified the paxillin LIM2 and LIM3 domains as being critical for focal adhesion targeting (Brown et al., 1996). Moreover, paxillin can localize to FAs in the absence of kindlins (Klapproth et al., 2019; Theodosiou et al., 2016) suggesting additional, kindlin-independent mode(s) of integrin engagement, presumably involving paxillin LIM2 and LIM3.

Here, we demonstrate that the paxillin LIM2 and LIM3 domains directly interact with carboxy-terminal residues of the integrin β subunit. Biochemical analysis of recombinant proteins, the NMR-based 3D-structure of the paxillin LIM domains, and functional analysis of mutated paxillin and integrin β3 in vitro and in the cellular context reveal that this interaction is based on a clamp-like extension in paxillin’s LIM3 domain, which contributes to cellular responses towards integrin ligands.

## Results

### Paxillin LIM2/3 domains can directly bind the cytoplasmic tails of integrin β1 and β3

In line with previous reports (Pasapera et al., 2010; Schiller et al., 2011), we recently observed that paxillin can be recruited in the absence of force to clusters of the integrin β CT, similar to the behaviour of known integrin binding partners such as talin and kindlin2 (Baade et al., 2019). We wondered whether recruitment to clustered integrin β tails is a general feature of LIM domain containing adhesome proteins. To this end, we used CEACAM-integrin β CT fusion proteins (CEA-ITGB1 or CEA-ITGB3), which can be engaged from the outside of the cell by multivalent CEACAM binding bacteria (*Neisseria gonorrhoeae*). This process initiates microscale accumulation of free integrin β tails mimicking nascent adhesion formation and was therefore named Opa protein triggered integrin clustering (OPTIC) (Suppl. Fig. S1A) (Baade et al., 2019). Interestingly, when co-expressed with CEA3-ITGB1 or CEA3-ITGB3, only paxillin and the closely related proteins Hic-5 and leupaxin showed a significant enrichment (Suppl. Fig. S1B-C). All other LIM domain proteins did not accumulate at clustered integrin β tails (Suppl. Fig. 1C-D). Paxillin recruitment to integrin clusters was dependent on the LIM domains, since the paxillin C-terminus encompassing LIM1-LIM4, but not the isolated N-terminal LD1-LD5 domains, strongly accumulated at integrin β cytoplasmic tails, and paxillin LIM1-LIM4, but not the LD1-5 domains, displaced full length paxillin from focal adhesion sites (Fig. 1A and B). These results suggest that paxillin, leupaxin and Hic-5 differ from other LIM domain containing adhesome proteins by their ability to locate at clustered integrin β tails and confirm the important role of the paxillin LIM domains for the specific subcellular localization. Moreover, pull-down assays with purified, recombinant proteins demonstrated that, similar to the talin F3 domain and Kindlin2, the paxillin LIM2/3 domains mediate a direct interaction of this adapter protein with the cytoplasmic tails of integrin β1 and β3 (Fig. 1C).

**Figure 1:**
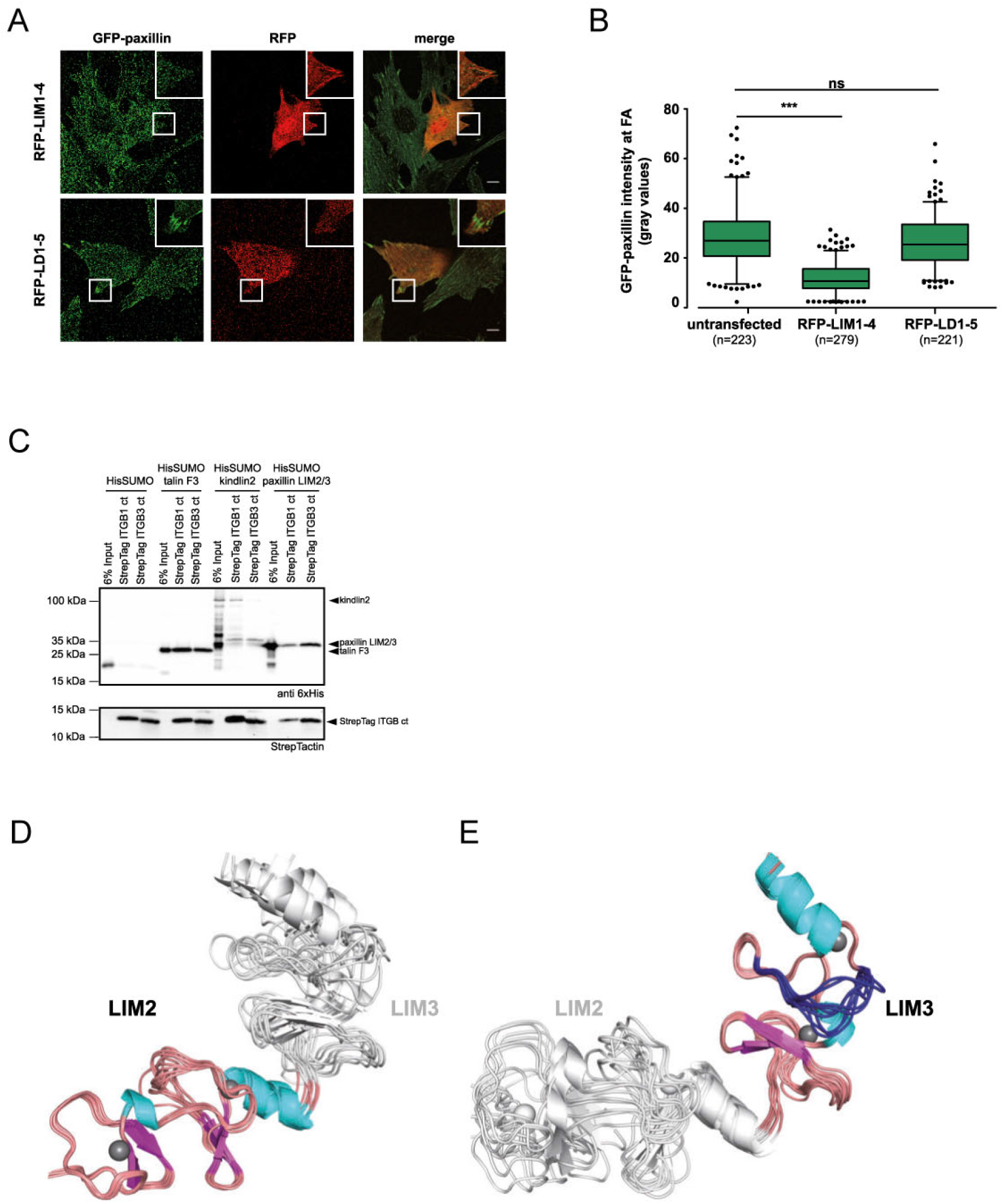
Paxillin LIM2/3 domains can directly bind the cytoplasmic tails of integrin β1 and β3. (A) Stable GFP-Paxillin expressing Flp-In 3T3 cells were transiently transfected with RFP-tagged LIM1-4 or LD1-5 domains. (B) Paxillin localization at FAs was evaluated by measuring the GFP fluorescence intensity in presence of either overexpressed LIM or LD domains. Shown are mean values of GFP-intensity from three independent experiments. The total number of analysed FAs is given in brackets under each sample. Error bars represent 5 and 95 percentiles. Significance was calculated using one-way ANOVA, followed by Bonferroni Multiple Comparison Test (*** p<0.0001, ns = not significant). (C) In vitro pulldown using recombinant Twin-Strep-tag integrin β cytoplasmic tails and recombinant talin1 F3 domain, full length kindlin2 as well as paxillin LIM2/3 domain fused to His_6_-Sumo, or His_6_-Sumo only as negative control. (D) and (E) Solution structure of paxillin LIM2/3. The final ensemble of ten conformers with lowest target function is shown fitted to the LIM2 domain (residues P381 to F438) (D) and fitted to the LIM3-domain (residues P440 to R497) (E), shown in ribbon representations. In the fitted part, α-helices are colored cyan and β-strands magenta. Zinc ions are shown as grey spheres. The flexible loop of the LIM3 domain (residue T473 to E482) is shown in blue. The domain that was not used for fitting is shown in light grey.

### The solution structure of paxillin LIM2/3 reveals a flexible loop region in the LIM3 domain

To investigate the direct interaction between paxillin and the integrin β subunit in more detail, we used NMR spectroscopy to gain structural insight and to delineate the binding interface. Since LIM3 and to a lesser extent LIM2 have been shown to be mainly responsible for FA targeting of paxillin (Brown et al., 1996), we expressed the LIM2/3 tandem domain (aa380-499) of human paxillin and determined its solution structure based on heteronuclear multidimensional NMR experiments (Sattler et al., 1999). Both the LIM2 and the LIM3 domain of paxillin exhibit the characteristic double zinc finger motif as described for other LIM domain containing proteins and paxillin family members (Kadrmas and Beckerle, 2004; Kontaxis et al., 1998; Matthews et al., 2009; Perez-Alvarado et al., 1994). Each domain comprises two orthogonally packed β-hairpins, followed by an α-helix (Fig. 1D and E). Interestingly, the LIM2 and LIM3 domains are connected by a short linker of 4 amino acids (F438-K441) that provides some degree of flexibility between both domains. This would fit into the previously proposed scenario of the LIM domains as a sort of molecular ruler and/or tension sensor (Schiller et al., 2011).

According to the measurements of the heteronuclear NOE between HN and N of the amides, the linker between LIM2 and LIM3 shows only slightly higher flexibility on the ps-to-ns timescale than the domains themselves (Fig. S2A). But there are less long-range NOE contacts in this region, suggesting that the structure is less densely packed here, resulting in differential relative orientations of the domains (Fig. 1D and E).

Interestingly, by analyzing the heteronuclear NOEs and the structural definition of the final ensemble of conformers, we identified a 7 amino acid stretch (F475-F481) in the second zinc-finger of the LIM3 domain that constitutes a flexible, surface exposed loop in the free protein (Fig. 1E). This loop is flanked by F475 and F481 and is situated adjacent to a hydrophobic patch or groove. In this region, the amino acid sequence and in particular residues F475, F480 and F481 are highly conserved within paxillin family members and across species (Suppl. Fig. S2B). We speculated that this flexible loop might act in concert with its opposing residues (E451, N452 and Y453) of the first zinc-finger and the hydrophobic patch to support a clamp-like mechanism for integrin β CT binding.

### The C-terminal residues of integrin β3 cytoplasmic tail are crucial for paxillin binding

To identify the binding region of paxillin LIM2/3 on the integrin β CT, we titrated unlabeled paxillin LIM2/3 to ^15^N labeled cytoplasmic tails of human integrin β1 (aa758-798) or β3 (aa748-788), respectively. In both cases, significant chemical shift perturbations (CSPs) of specific integrin residues could be observed (Fig. 2A and B, and Suppl. Fig. S3A and B). A dissociation constant (K_D_) of 52±30 µM could be determined for integrin β1 (Suppl. Fig. S3A), while integrin β3 showed a higher K_D_ (528±130 µM) (Fig. 2A). These findings are in line with our previous microscopic observations, where recruitment of paxillin to CEA-ITGB1 was more pronounced (Suppl. Fig. 1C). Surprisingly, when mapping the CSPs onto the primary sequence of integrin β CTs, the interacting regions were distinct. While the highest CSPs in integrin β1 CT were distributed over the membrane proximal NPxY motif and a neighbouring conserved TT motif (Suppl. Fig. S3B), the largest chemical shifts in integrin β3 were confined to the eight C-terminal amino acids, spanning the membrane distal NxxY motif (Fig. 2B). Indeed, deleting eight amino acids from the C-terminus of integrin β3 (Δ8aa) completely abrogated paxillin LIM2/3 binding in pulldown experiments, while deletion of the last three amino acids (Δ3aa) significantly reduced paxillin binding (Fig. 2C). As expected, applying either of these mutant integrin β3 CTs in titration experiments with paxillin LIM2/3 yielded no significant CSPs, confirming the loss of interaction (Fig. 2D and E and Suppl. Fig. S3C-D). The biochemical data were further corroborated by OPTIC assays, were both the Δ3aa and the Δ8aa mutant, but not the S778A mutation in the kindlin binding site, diminished paxillin recruitment to clustered integrin β3 tails (Suppl. Fig. S3E-F).

**Figure 2:**
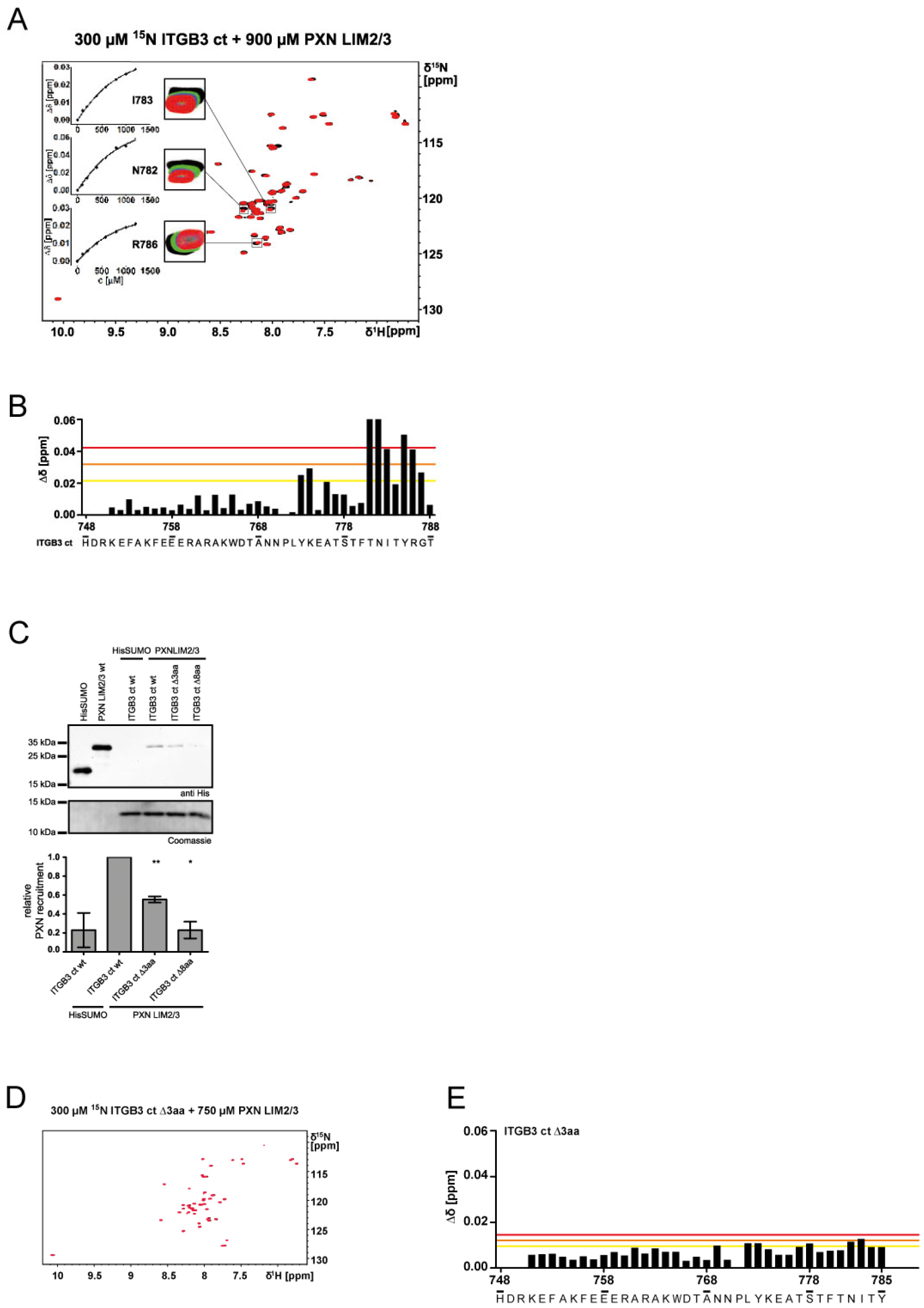
The C-terminal residues of integrin β3 cytoplasmic tail are crucial for paxillin binding. (A) ^15^N-HSQC titration of 300 µM ^15^N integrin β3 ct (ITGB3 ct) with paxillin LIM2/3. Paxillin was added in concentrations up to 900 µM. Boxes show a selection of signals affected by CSPs (residues N782, I783 & R786) in the presence of 0 µM (black), 150 µM (green), 300 µM (blue) and 900 µM (red) paxillin LIM2/3. Insets show the concentration dependence of combined amide CSPs globally fitted to a one site binding model. (B) Combined amide CSPs of 300 µM ^15^N integrin β3 ct in the presence of 760 µM paxillin LIM2/3 vs residue number of integrin β3 ct. Lines indicate average δΔ + 1x s.d. (yellow), δΔ + 2x s.d. (orange) and δΔ + 3x s.d. (red). (C) Representative Western Blot of in vitro pulldown using biotinylated integrin β peptides and recombinant His_6_-SUMO or His_6_-SUMO-paxillin LIM2/3 (PXN LIM2/3). A decreased binding of paxillin to truncated integrin β3 peptides is visible. Lower panel: Densitometric quantification of the Western Blots in (A) (n=3). Statistical significance was calculated using One sample t-test to calculate if samples mean are significantly different from a hypothetical value of 1 (* p<0.05, ** p< 0.01). (D) ^15^N-HSQC titration of 300 µM ^15^N integrin β3 ∆3aa (ITGB3 ∆3aa) with paxillin LIM2/3 (PXN LIM2/3). Paxillin was added up to a concentration of 750 µM. (E) Combined amide CSPs of 300 µM ^15^N integrin β3 ct ∆3aa in the presence of 750 µM paxillin LIM2/3 vs residue number of integrin β3 ct ∆3aa.

### Paxillin directly associates with the C-terminus of integrin β3 to contribute to cell spreading

To study the physiological relevance of these mutations, we generated integrin β3 knockout fibroblasts via CRISPR/Cas9 and stably re-introduced either full-length integrin β3 wt or one of the truncated integrin β3 variants, Δ8aa or Δ3aa (Suppl. Fig. S4A and B). While integrin β3-deficient cells exhibited a strongly impaired initial spreading on vitronectin- and fibronectin-coated substrates, the re-expression of integrin β3 wt reverted this phenotype (Fig. 3A and B). In contrast, integrin β3 Δ3aa, and even more so integrin β3 Δ8aa re-expressing fibroblasts were still impaired in their spreading ability (Fig. 3A and B). A similar spreading defect on integrin ligands has also been reported for kindlin-deficient cells and deletions of the integrin carboxy-terminus might also corrupt the kindlin binding site, indirectly affecting the recruitment of paxillin. To substantiate our biochemical findings of a direct paxillin interaction with the integrin β subunit, we employed kindlin1 and 2-deficient double knock-out cells (Bottcher et al., 2017; Theodosiou et al., 2016). As these cells display strongly diminished expression of integrin β3, we introduced integrin β3 wt, integrin β3 Δ8aa, or integrin β3 Δ3aa into the kindlin1/2 KO cells (Suppl. Fig. S4C and D). As reported before (Bottcher et al., 2017), the kindlin1/2 deficient fibroblasts hardly attached and did not spread on the vitronectin-coated substrate (Fig. 3C). However, paxillin-positive focal attachment sites appeared upon expression of integrin β3 wt in the kindin1/2 deficient cells, whereas no such matrix contact sites were detectable in cells expressing the truncated integrin β3 mutants (Fig. 3C). The adhesions appeared upon plating of the integrin β3 wt cells onto vitronectin, but not upon plating onto poly-L-lysine (Fig. 3C and Suppl. Fig. S4E), and these integrin β3-mediated contacts also stained positive for talin (Fig. 3D and Suppl. Fig. S4E). Altogether, our results demonstrate that paxillin can localize to FAs in the absence of kindlins and that this process requires the paxillin binding site in the integrin β3 cytoplasmic tail.

**Figure 3:**
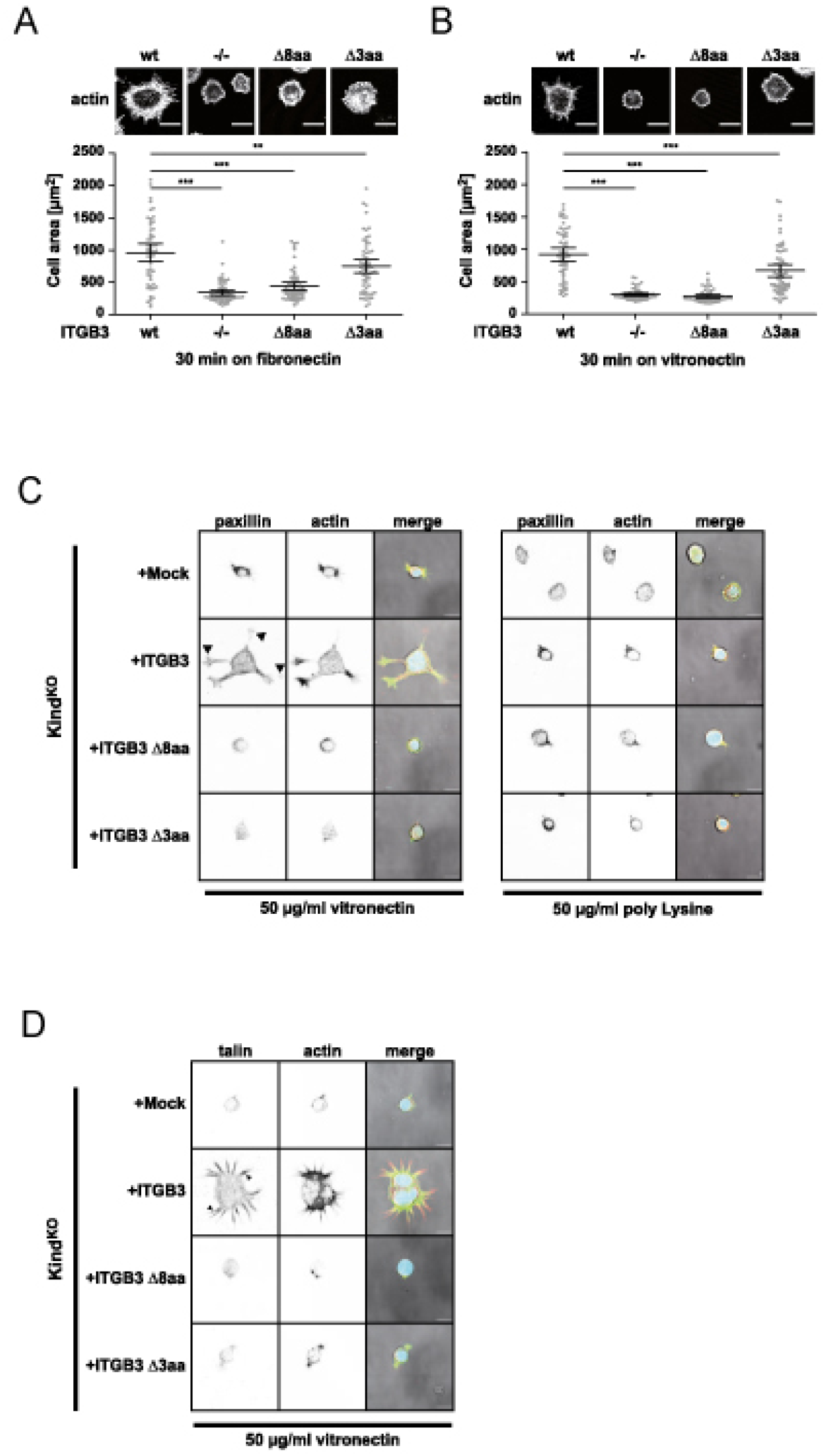
C-terminally truncated integrin β3 causes defective cell spreading. (A) and (B) Serum starved Flp-In 3T3 integrin β3 knockout fibroblasts or knockout cells re-expressing integrin β3 wt or C-terminally truncated integrin β3 mutants were seeded onto glass slides coated with 5 µg/ml fibronectin (A) or 5 µg/ml vitronectin (B) for 30 min and cell area was measured. Shown are mean values and 95% confidence intervals of n=60 cells per sample from 3 independent experiments. Statistical significance was calculated using one-way ANOVA followed by Bonferroni Multiple Comparison Test. (C) Serum starved kindlin1/2 deficient fibroblasts (Kind^KO^) cells stably expressing full length integrin β3 or truncated mutants were seeded on glass coverslips coated with 50 µg/ml vitronectin (left panel) or poly-Lysin (right panel) for 4 h. Cells were fixed and stained for endogenous paxillin. (D) Serum starved Kind^KO^ cells stably expressing full length integrin β3 or truncated mutants were seeded on glass coverslips coated with 50 µg/ml vitronectin for 4 h. Cells were fixed and stained for endogenous talin.

### The LIM3 flexible loop mediates direct association with the integrin β3 cytoplasmic tail

To precisely identify the integrin binding site within the paxillin LIM2/3 domains, we titrated the unlabeled CT of human integrin β3 to ^15^N-labelled paxillin LIM2/3. Similar to the inverse titrations, the K_D_ value of this interaction was determined to 532±239 µM (Suppl. Fig. S5A). Importantly, the most prominent CSPs in paxillin were recorded within the second zinc-finger of the LIM3 domain, specifically in the flexible loop region between F475 and F481 (Fig. 4A, B and C). To verify our NMR based epitope mapping we individually mutated residues of the loop region. Interestingly, exchanging phenylalanine F475, F480, or F481 for alanine caused a complete or partial unfolding of the LIM3 domain (Suppl. Fig. S6A-F). Though these residues show strong CSPs and participate in integrin binding, these phenylalanines are also essential for maintaining the structure of the LIM domain. To gain further insight into the role of the loop region, these phenylalanines were left intact, but instead the residues between F475 and F480 were mutated to alanine (LIM2/3-4A). Paxillin LIM2/3-4A exhibited a stable LIM domain fold, however, no saturable CSPs could be detected when titrating up to millimolar concentrations of integrin β3 to ^15^N-labelled LIM2/3-4A (Suppl. Fig. S7A and B). Pulldown assays with the paxillin LIM2/3-4A mutant confirmed diminished binding to the integrin β3 CT (Fig. 4D). Next, we introduced the LIM3 loop mutations into full length paxillin (GFP-PXN-4A) to corrupt the direct engagement of integrin β3 in intact cells. In addition, we generated a truncated paxillin lacking the LIM4 domain (GFP-PXN ΔLIM4) to interfere with the kindlin-mediated indirect binding of paxillin to the integrin beta subunit. GFP-tagged paxillin wt, GFP-PXN-4A, or GFP-PXN ΔLIM4, were re-introduced into paxillin knockout fibroblasts and their cell spreading was monitored (Fig. 4E and F). As expected, spreading of paxillin KO cells on vitronectin was strongly impaired. Re-expression of GFP-paxillin wt rescued this phenotype, whereas the GFP-PXN-4A mutant was not able to revert the spreading defect and mimicked paxillin KO cells in the first 2 h after seeding on the substrate (Fig. 4E and F). In contrast, paxillin lacking the LIM4 domain reconstituted cell spreading in a similar manner as wildtype paxillin (Fig. 4E and F). Moreover, paxillin wildtype and PXN ΔLIM4 reverted the round, circular morphology of the paxillin-knock-out fibroblasts to the spindle shaped, pointed cell phenotype of the wildtype fibroblasts, whereas cells re-expressing GFP-PXN-4A retained the elevated circularity of the paxillin knock-out cells (Fig. 4E). Taken together, these results underscore the functional importance of the direct interaction between paxillin and integrin β3. The clamp-like mechanism afforded by the flexible loop in paxillin’s LIM3 domain could stabilize the association with the integrin β3 cytoplasmic domain and modulates integrin β3-initiated cellular responses.

**Figure 4:**
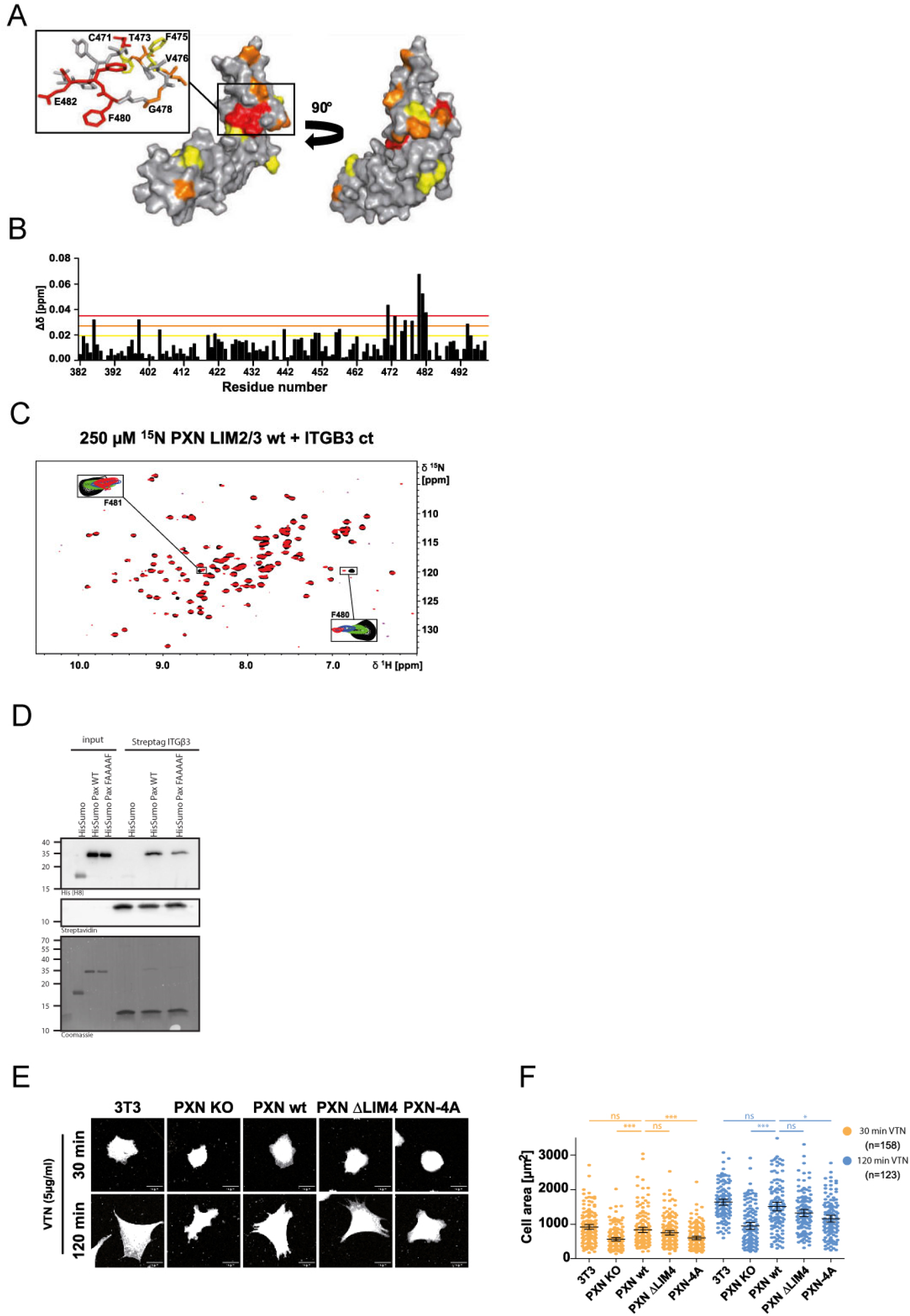
The LIM3 flexible loop mediates binding to the integrin β cytoplasmic tail. (A) Mapping of combined amide CSPs when binding to integrin β3 onto the solution structure of paxillin LIM2/3 shown as surface representation from two perspectives. Residues showing CSPs larger than average δΔ + 3x s.d. are colored red, residues for which [average δΔ + 3x s.d. < δΔ < average δΔ +2x s.d.] are colored orange, and residues for which [average δΔ + 2x s.d.< δΔ < average δΔ + 1x s.d.] are colored yellow. Residues with δΔ < average + 1x s.d. are colored grey. The boxed region is also shown in stick representation including the flexible loop of the LIM3 domain using the same color code. Residues experiencing significant CSPs are labelled by amino acid type and position. (B) Combined amide CSPs of 250 µM ^15^N paxillin LIM2/3 in the presence of 750 µM integrin β3 vs residue number of paxillin. Lines indicate average δΔ + 1x s.d. (yellow), δΔ + 2x s.d. (orange) and δΔ + 3x s.d. (red). (C) ^15^N-HSQC titration of 250 µM ^15^N paxillin LIM2/3 wt (PXN LIM2/3 wt) with integrin β3. Integrin was added in concentrations up to 2420 µM. Boxes show a selection of signals affected by CSPs (residues F480 & F481) in the presence of 0 µM (black), 200 µM (green), 600 µM (blue) and 2420 µM (red) integrin β3 ct. (D) Recombinantly expressed His_6_-SUMO PXN LIM2/3 wt or His_6_-SUMO PXN LIM2/3 4A were pulled down using Twin-Strep-Tag integrin β3 ct. PXN LIM2/3 4A shows reduced binding to integrin β3 ct. (E) Outside-in signalling dependent cell spreading of paxillin knockout cells, stably re-expressing ctrl vector (PXN KO), GFP-paxillin wt (PXN wt) or paxillin mutants PXN ∆LIM4 and PXN-4A. Cells were starved overnight and seeded for 30 or 120 min, respectively on the integrin β3 ligand vitronectin in the absence of serum. Cells were fixed and the cell membrane was stained with CellMask Orange. Scale bar represents 20 µm. (F) Quantification of cell area from images (E). Shown are mean values with 95% confidence intervals from 3 independent experiments for 30min timepoint or from 2 independent experiments for 120 min timepoint. Sample sizes are given in brackets. Statistical significance was calculated using one-way ANOVA followed by Bonferroni Multiple Comparison Test (ns: not significant; *** p≤0.0001; ** p≤0.01).

## Discussion

Although paxillin was discovered more than 30 years ago and constitutes a core focal adhesion protein, its mode of focal adhesion targeting has remained controversial. Recent biochemical approaches have pointed to an indirect association of paxillin with the integrin β1 and β3 subunits via the integrin binding partners kindlin1 or kindlin2 (Gao et al., 2017; Theodosiou et al., 2016). As the association of kindlin with paxillin appears to rest on the paxillin LD repeats and the LIM4 domain (Bottcher et al., 2017; Zhu et al., 2019), these findings do not explain the central role of the LIM3 domain for focal adhesion localization, which has been delineated by microscopic observations in intact cells (Brown et al., 1996). Here, we present evidence for a direct interaction between the paxillin LIM3 domain and the cytoplasmic tails of integrin β1 and integrin β3, respectively. Together with the indirect link provided by kindlin, the intimate association of paxillin with the integrin β subunit now unveils the full spectrum of paxillin’s focal adhesion recruitment modalities and unmasks the fundamental building principles of cellular attachment sites.

In solution, paxillin’s LIM2 and LIM3 domains adopt an overall fold consistent with available structural data for LIM domains of other proteins (Kadrmas and Beckerle, 2004; Kontaxis et al., 1998; Matthews et al., 2009; Perez-Alvarado et al., 1994). However, our NMR structure reveals an intriguing detail, which is conserved in paxillin orthologues from other species. Indeed, the LIM3 domain of paxillin harbours a flexible, surface exposed loop, which, based on sequence homology, is also present in the paxillin family members Hic-5 and leupaxin. This loop demarcates the integrin binding site in the LIM3 domain and appears to function as a clasp to stabilize the association of paxillin with the integrin cytoplasmic tail. This additional direct interaction between the integrin β cytoplasmic tail and paxillin now consolidates an emerging principle of focal adhesion organization: each core component of focal adhesions, including talin, kindlin, paxillin, FAK, vinculin, and α-actinin is able to sustain multiple, independent interactions with other FA components. In analogy to a steel frame construction, this kind of assembly not only allows a stepwise expansion of the protein complex, but also provides a further mechanical reinforcement with every incoming component. In the specific example of paxillin, this protein can associate via its LIM4 domain with integrin-bound kindlin2 (Zhu et al., 2019), but then paxillin will also be in place to exploit the clamping mechanism build in its LIM3 domain to bind the integrin β subunit and to re-enforce this tripartite complex as a pre-requisite for efficient initial cell spreading. Furthermore, as paxillin can interact via its LIM1/2 domains with the talin head (Ripamonti et al., 2021) and via its LD1 domain with the talin rod region (Zacharchenko et al., 2016), there is the possibility that paxillin strengthens the early integrin-associated protein complex beyond kindlin. Indeed, a stabilization of the integrin-talin-kindlin nexus by paxillin has been observed (Gao et al., 2017). Multiple reciprocal interactions between talin, kindlin, paxillin and the integrin β tail would also increase the avidity and might demonstrate once more how multivalent low-affinity interactions such as those observed for paxillin LIM3 and integrin β3 stabilize macromolecular networks. Indeed, the relatively low affinity between LIM3 and integrin β3 might be a prerequisite for efficient assembly and disassembly of FAs.

Intriguingly, such reciprocal interactions between focal adhesion core components could also be the basis for the astonishing flexibility in the temporal sequence, in which these proteins can assemble at integrin cytoplasmic domains. For example, talin binding to the integrin β tail appears as a pre-requisite for the recruitment of its binding partner FAK (Wang et al., 2011; Zhang et al., 2008), while the opposite sequence of assembly has also been reported (Lawson et al., 2012; Lawson and Schlaepfer, 2012). This behaviour is mirrored by kindlin and paxillin, as kindlin is able to recruit paxillin, while the direct binding of paxillin to the β subunit can turn this sequence of events on its head and could potentially allow paxillin-dependent recruitment of kindlin. Interestingly, one of the factors, which determines the order of assembly appears to be the nature of the involved integrin heterodimer. In particular, differences between integrin α5β1 and integrin αvβ3 do not only exist with regard to the ligand spectrum, but also how they convey the ligand binding event into the cell (Bachmann et al., 2019). In this regard, α5β1 integrins are known to determine adhesion strength and form catch bonds with their ligands, when increasing forces are applied (Kong et al., 2009; Roca-Cusachs et al., 2009). The elevated binding affinity of paxillin for integrin β1 and its ability to associate with talin and kindlin might reflect this need to withstand high forces. Though integrin αvβ3 is not able to sustain the high binding strength of α5β1 integrin, integrin αvβ3 exhibits faster binding rates and stimulates integrin α5β1-mediated binding (Bharadwaj et al., 2017; Schiller et al., 2013). Interestingly, the fast binding rate of integrin αvβ3 correlates with its increased ability to recruit paxillin and its propensity to initiate larger paxillin-positive adhesion sites (Bharadwaj et al., 2017; Missirlis et al., 2016). Furthermore, on patterned substrates, cell spreading and paxillin recruitment preferentially occur via the vitronectin-binding integrin αvβ3 (Pinon et al., 2014). Together with our findings of a prominent recruitment of paxillin to integrin β3 in kindlin-deficient cells and of reduced spreading of paxillin LIM3-4A expressing cells on vitronectin, all these observations suggest a particularly prominent role for the direct association of the paxillin LIM3 domain with integrin β3. It is also interesting to note that the paxillin LIM3 domain appears to latch onto a specific section of the integrin β3 subunit at the far carboxy-terminus, which shows significant chemical shift perturbations upon binding. The C-terminal amino acids of integrin β3 differ from all other β subunits making this a unique recognition site and helping to explain this peculiar interaction mode of paxillin and integrin β3.

Though further studies are needed to delineate the stoichiometry of talin, kindlin and paxillin at clustered integrin β subunits, our structural elucidation of the paxillin LIM2 and LIM3 domains and their association with the integrin β carboxy-terminus now provides the foundation to probe and manipulate the functional contribution of paxillin to matrix adhesion and cell spreading.

## Material & Methods

### Antibodies and dyes

The following primary and secondary antibodies were used at indicated concentrations: anti-human α-actinin1 (mouse monoclonal, BM75.2, Sigma Aldrich, A5044; WB 1:1000), anti-human talin (mouse monoclonal, 8d4, Sigma Aldrich, T3287; WB 1:750), anti-human FAK (rabbit polyclonal, A-17, Santa Cruz, sc-557; WB 1:250), anti-human kindlin2 (mouse monoclonal, 3A3, Merck, MAB2617; WB 1:1000, IF 1:200), anti-mouse kindlin2 (rabbit polyclonal, 11453-1-AP, Proteintech; WB 1:2000), anti-human cSRC (rabbit polyclonal, SRC2, Santa Cruz, sc-18; WB 1:1000), anti-human ILK (rabbit monoclonal, EP1593Y, Epitomics; WB 1:1000), anti-human p130Cas (rabbit polyclonal, N17, Santa Cruz; WB 1:1000), anti-human vinculin (mouse monoclonal, VIN-1, Sigma Aldrich, V9131; WB 1:1000), anti-human Hic-5 (mouse monoclonal, 34, BD Biosciences, 611164; WB 1:500), anti-human paxillin (mouse monoclonal, 5H11, Thermo Fisher Scientific, AHO0492; WB 1:1000, IF 1:200), anti-GAPDH (mouse monoclonal, GA1R, Thermo Fisher Scientific, MA5-15738-HRP), anti-human Rac (rabbit polyclonal, invitrogen, PA5-17519; WB 1:1000), anti-human CEACAM1, 3, 4, 5, 6 (mouse monoclonal, D14HD11, Aldevron; WB 1:6000, IF 1:200); mouse monoclonal anti 6xHis (mouse monoclonal, HIS.H8, Thermo Fisher Scientific, MA1-21315; WB 1:2000), anti GFP (mouse monoclonal, JL8, Clontech; WB: 1:6000), anti-human tubulin (mouse monoclonal, E7, purified from hybridoma cell supernatants, Developmental Studies Hybridoma Bank, University of Iowa, USA; WB 1:1000), anti-mouse integrin β1 (Armenian hamster monoclonal, Hmb1-1, Thermo Fisher Scientific, 11-0291-82; FC 1:300), anti-mouse integrin β3 (armenian hamster monoclonal, 2C9.G3, Thermo Fisher Scientific, 13-0611-81; FC 1:200), anti-mouse integrin α5 (rat monoclonal, MFR5, BD Biosciences, 553319; FC 1:300), anti-mouse integrin αV (rat monoclonal, RMV-7, BD Biosciences, 550024; FC 1:300). Secondary antibodies used: horseradish peroxidase (HRP)-conjugated goat anti-mouse; WB: 1:10.000, horseradish peroxidase (HRP)-conjugated goat anti-rabbit; WB: 1:5000; Cy5-conjugated goat anti-mouse; IF 1:200, Dylight 488 conjugated goat anti-mouse, IF 1:200, Rhodamine Red-X conjugated goat anti-arm. Hamster; FC 1:300, Rhodamine Red conjugated goat anti-rat; FC 1:300 (all from Jackson ImmunoResearch Inc., Baltimore, USA). CellMask Orange Plasma membrane stain, Thermo Fisher Scientific, C10045; IF 1:1000.

### Cell culture and transient transfection

Human embryonic kidney 293T cells (293T; American Type Culture Collection CRL-3216) were grown in DMEM supplemented with 10% calf serum. Flp-In™ 3T3 cells (Thermo Fisher Scientific) were cultured in DMEM supplemented with 10% fetal calf serum (FCS) and 1% non-essential amino acids. GFP-FAK expressing mouse embryonic fibroblasts (GFP-FAK MEFs) derived from FAK/p53 -/- knockout MEFs (Schlaepfer et al., 2007) were cultured in DMEM supplemented with 10% FCS and 1% non-essential amino acids on gelatine-coated (0.1% in PBS) cell culture dishes. All cells were maintained at 37°C, 5% CO_2_, and subcultured every 2–3 days.

For transient transfection of 293T cells, cells were seeded at 25% confluence the day before and transfected using the standard calcium phosphate method with a total amount of 5 µg plasmid DNA/dish. For transient transfection of Flp-In™ 3T3 cells, 1 × 10^5 cells were seeded into 6 well plates the day before and transfected with using jetPRIME® transfection reagent (Polyplus transfection, Illkirch, France), following manufacturer’s protocol. GFP-FAK MEFs were transiently transfected with Lipofectamin 2000, according to manufacturer’s recommendations.

### Whole cell lysates (WCLs) and WB

WCLs were generated by lysing equal cell numbers in radioimmunoprecipitation assay buffer (1% Triton X-100, 50 mM Hepes, 150 mM NaCl, 10% glycerol, 1.5 mM MgCl_2_, 1 mM EGTA, 0.1% wt/vol SDS, and 1% vol/vol deoxycholic acid) supplemented with freshly added protease and phosphatase inhibitors (10 mM sodium pyrophosphate, 100 mM NaF, 1 mM sodium orthovanadate, 5 µg/ml leupeptin, 10 µg/ml aprotinin, 10 µg/ml Pefabloc, 5 µg/ml pepstatin, and 10 µM benzamidine) and phosphatase saturating substrate (para-nitrophenolphosphate [pNPP]; Sigma-Aldrich; 10 mM). Chromosomal DNA was mechanically sheared by passing through a metal needle. DNA and cell debris were pelleted by addition of sepharose beads and centrifugation (13,000 rpm, 30 min, 4°C). Supernatant was supplemented with 4× SDS sample buffer (4% wt/vol SDS, 20% wt/vol glycerol, 125 mM Tris-HCl, 20% vol/vol β-mercaptoethanol, and 1% wt/vol Bromophenol blue, pH 6.8) and boiled for 5 min at 95°C. Proteins were resolved on 10–18% SDS-PAGE. After separation, the proteins were transferred to a polyvinylidene fluoride membrane (Merck Millipore), followed by blocking in 2% BSA containing 50 mM Tris-HCl, 150 mM NaCl, and 0.05% Tween 20, pH 7.5 (TBS-T) buffer. The membrane was incubated with primary antibody in blocking buffer overnight at 4°C, washed three times with TBS-T, and incubated with HRP-conjugated secondary antibody in TBS-T for 1 h at RT. The chemiluminescent signal of each blot was detected with ECL substrate (Thermo Fisher Scientific) on the Chemidoc Touch Imaging System (Bio-Rad) in signal accumulation mode. Acquired images were processed in Adobe Photoshop CS4 by adjusting illumination levels of the whole image.

### Recombinant DNA

The construction of His_6_SUMO-tagged talin F3, eGFP-tagged full length talin, His_6_SUMO-kindlin2 as well as the generation of the Twin-Strep-tag vector for bacterial expression has been described in detail (Grimm et al., 2020). The generation of CEA3-ITGBct fusion constructs has been described previously (Baade et al., 2019). cDNA of human paxillin isoform a (NM_002859.4) was kindly provided by Alexander Bershadsky (Mechanobiology Institute, National University of Singapore, Singapore) and was used as template for PCR amplification. His_6_SUMO-PXN LIM2/3 was generated by amplifying paxillin using primers: PXN LIM2/3 forward: 5’-CCAGTGGGTCTCAGGTGGTTCCCCGCGCTGCTAC-3’; PXN LIM2/3 reverse: 5’-CTGATCCTCGAGTTACCCATTCTTGAAATATTCAGGCGAGCCGCGCCGCTC-3’.

The product was then ligated into pET24a His-Sumo bacterial expression vector using Eco31I and XhoI restriction sites.

Paxillin full length was amplified using primers PXN-fl forward: 5’-ACTCCTCCCCCGCCATGGACGACCTCGACGCCCTGCTG-3’ and PXN-fl reverse: 5’-CCCCACTAACCCGCTAGCAGAAGAGCTTGAGGAAGCAGTTCTGACAGTAAGG-3’. Paxillin LD1-5 was amplified using primers PXN-fl forward and PXN LD1-5 reverse: 5’-CCCCACTAACCCGCAGCTTGTTCAGGTCAG-3’. Paxillin LIM1-4 was amplified using primers PXN-LIM1-4 forward: 5’-ACTCCTCCCCCGCCATGAAGCTGGGGGTCGCCACAGTCGCCAAAG-3’ and PXN-fl reverse.

cDNAs encoding LIM domain proteins: Hic-5 cDNA (isoform 1, NM_001042454.3) was kindly provided by Nicole Brimer (University of Virginia, Charlottesville, USA). Hic-5 was amplified using the primers Hic-5 forward: 5’-ACTCCCCCGCCATGGAGGACCTGGATGCCC-3’ and Hic-5 reverse: 5’-CCCCACTAACCCGTCAGCCGAAGAGCTTCAGG-3’. Leupaxin was amplified from pOTB7-LPXN (obtained from Harvard Medical School PlasmID Database; HsCD00331641) using primers LPXN forward: 5’-ACTCCTCCCCCGCCATGGAAGAGTTAGATGCC-3’ and LPXN reverse: 5’-CCCCACTAACCCGGCATTACAGTGGGAAGAGC-3’. PINCH-1 was amplified from pDNR-LIB hLIMS1 (obtained from Harvard Medical School PlasmID Database, HsCD00326503) using the primers PINCH forward: 5’-ACTCCTCCCCCGCCATGGCCAACGCCCTGGCCAGC-3’ and PINCH reverse: 5’-CCCCACTAACCCGTTTCCTTCCTAAGGTCTCAGC-3’. cDNA for LASP-1 was provided by Elke Butt (Universitätsklinikum Würzburg, Würzburg, Germany) and amplified using primers LASP forward: 5’-ACTCCTCCCCCGCCATGAACCCCAACTGCGCC-3’ and LASP reverse: 5’-CCCCACTAACCCGTCAGATGGCCTCCACGTAGTTGG-3’.

The respective PCR products were cloned into the pDNR-Dual-LIC vector according to the ligation independent cloning (LIC) strategy. The sequence verified constructs were then subcloned into the expression vector pEGFP-C1 harbouring a loxP recombination site via Cre-Lox recombination. mEmerald-Migfilin was a gift from Michael Davidson (Addgene #54182) and was used unmodified. pEGFP-Zyxin was described elsewhere (Agerer et al., 2005).

### Site directed mutagenesis

The amino-acid residue of interest was changed using the overlap-extension PCR mutagenesis procedure. Desalted oligonucleotide primers were purchased from Sigma-Aldrich (Merck KGaA, Darmstadt, Germany) and can be found in following table. Human Paxillin LIM2/3 was used as a template.

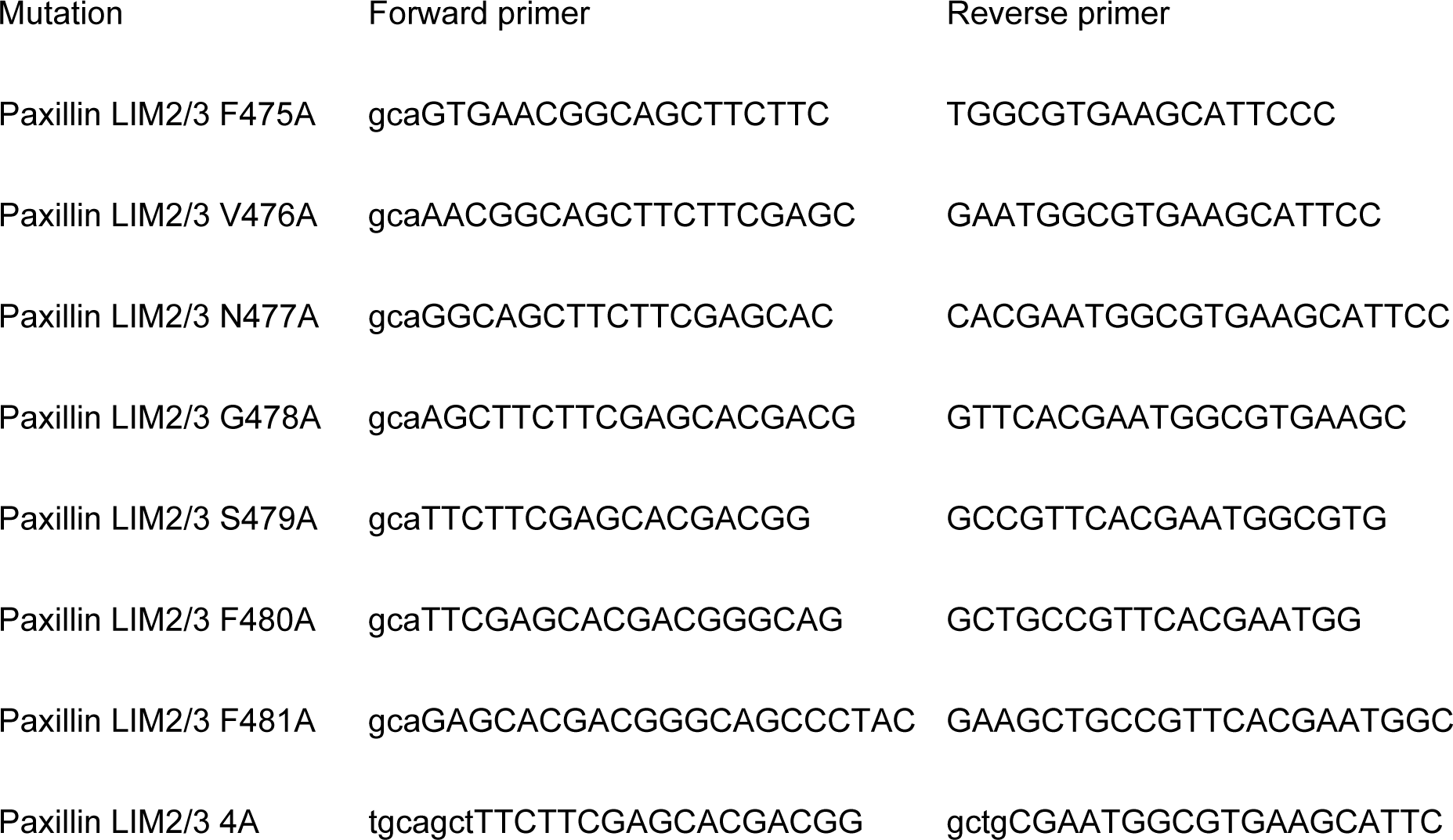

### Recombinant protein expression

Recombinant proteins with His_6_-SUMO tag encoded on a pET24a vector were expressed in *E. coli* Tuner (DE3) cells. Cells were cultured in lysogeny broth medium containing 50 µg/mL kanamycin and 1% glucose (wt/vol) (for paxillin constructs additionally 0.1 mM ZnCl_2_) at 37 °C until an OD_600_ value of 0.6-0.8 was reached. Subsequently, overexpression was induced by addition of isopropyl β-D-thiogalactoside (IPTG) to a final concentration of 0.5 mM (for integrin β1 1 mM respectively). After 6-8 h incubation (for integrin-β1 overnight, respectively) at 30 °C cells were harvested by centrifugation at 10 000 x g for 15 min at 4 °C and stored at - 80 °C. For isotopic labelling bacteria were cultured in M9-minimal medium containing ^15^N ammonium chloride and/or ^13^C glucose as sole nitrogen and carbon sources. Integrin cytoplasmic domains with a TwinStrepTagII tag encoded on a pET24a vector were expressed in *E. coli* BL21(DE3) pRosetta cells. Cells were cultured in lysogeny broth medium containing 50 µg/mL kanamycin. Expression conditions were identical to His_6_-SUMO integrin β cytoplasmic tails. His_6_-SUMO, His_6_-SUMO tagged talin F3 and kindlin2 were expressed in *E. coli* BL21(DE3). Bacteria were grown at 37 °C to an OD of 0.6-0.8 and induced with 1 mM IPTG overnight at 30°C (His_6_-SUMO and talin F3) or 20 °C (kindlin2).

### Protein purification

All steps were performed at 4 °C. Pelleted cells were slowly thawed on ice, resuspended in 1:5 (wt/vol) lysis buffer (50 mM Tris, 300 mM NaCl, pH 8.0, protease inhibitors) and lysed via a high-pressure homogenizer (Emulsiflex C3, Avestin Inc., Ottawa, Canada). The mixture was separated by ultracentrifugation at 100 000 x g for 30 min at 4 °C and supernatant was loaded onto a HisTrap HP column (GE Healthcare, Freiburg, Germany) preequilibrated with 50 mM Tris, 10 mM imidazole and 300 mM NaCl, pH 8.0. The loaded column was washed and eluted fractions, monitored by UV absorbency at 280 nm, were pooled and dialyzed in 50 mM Tris, 300 mM NaCl, pH 8.0. After overnight cleavage with Ulp1 the His_6_-SUMO tag was removed by subsequent HisTrap purification and protein solution was subjected to size-exclusion by using HiLoad 16/60 Superdex 30 (for integrin constructs) or Superdex 75 column (for paxillin constructs, GE Healthcare, Freiburg, Germany) preequilibrated with 50 mM Na_2_HPO_4_, 150 mM NaCl, pH 6.2 (for integrin constructs) or 7.5 (for paxillin constructs). For paxillin constructs all buffers contained also 0.1 mM ZnSO_4_ and 1 mM DTT respectively. Purified protein was checked by SDS-PAGE.

### Pulldown assays with integrin β cytoplasmic domains

2.5 μg of TwinStrep-tagged integrins or 10 μg of biotin-integrin peptides β3wt aa742-788 (Biotin-HDRKEFAKFEEERARAKWDTANNPLYKEATSTFTNITYRGT-OH), β3∆3aa aa742-785 (Biotin-HDRKEFAKFEEERARAKWDTANNPLYKEATSTFTNITY-OH) β3∆8aa aa742-780 (Biotin-HDRKEFAKFEEERARAKWDTANNPLYKEATSTFT-OH); all from Novopep Limited) were loaded onto Strep-Tactin Sepharose beads (50% suspension; IBA Lifesciences) or streptavidin agarose beads (50% suspension; 16-126; Merck) in pulldown buffer (50 mM Tris, pH 8, 150 mM NaCl, 10% glycerol, 0.05% Tween, 10 µM ZnCl_2_) for 30 min at RT under continuous rotation. After centrifugation (2,700 g, 2 min, 4°C), samples were washed three times with pulldown buffer. Then integrin-loaded beads were suspended in bait protein solution (2 µM of protein diluted in pulldown buffer) and incubated 2 h at 4°C under constant rotation. Samples were centrifuged (2,700 g, 2 min, 4°C) and washed three times with pulldown buffer. Strep-Tactin samples were eluted under native conditions by adding 30 μl of buffer BXT (50 mM Tris, pH 8, 150 mM NaCl, 50 mM biotin). After 10 min incubation at RT under constant rotation, samples were centrifuged. Supernatants were mixed with 4× SDS and boiled for 5 min at 95°C before they were subjected to WB. Streptavidin agarose beads were directly mixed with 2× SDS and boiled for 10 min at 95°C to elute proteins from biotin-integrin peptides before they were subjected to WB.

### Resonance assignment

All NMR-experiments for the resonance assignment and structure determination were recorded on a Bruker Avance III 600 MHz spectrometer equipped with an H/C/N TCI cryoprobe. Three-dimensional spectra were recorded using non-uniform sampling (25-50% sparse sampling) and reconstructed by recursive multidimensional decomposition (Topspin® v3.1-3.2). NMR-sample conditions: 500 μM ^13^C-^15^N-Paxillin-LIM2/3, 150 mM NaCl, 50 mM Na_2_HPO_4_, 4 mM NaN_3_, 1 mM DTT, 5% (or 100% D_2_O), pH 7.5. Recorded 3D-spectra in 5% D_2_O: HNCO, HN(CA)CO, CBCANH, CBCA(CO)NH, H(CCCO)NH, (H)C(CCO)NH, NOESY-^15^N-HSQC, NOESY-^13^C_ali._-HSQC; in 100% D_2_O: H(C)CH-TOCSY, (H)CCH-TOCSY, H(C)CH-COSY, NOESY-^13^C_ali._-HSQC, NOESY-^13^C_aro._-HSQC (NOESY mixing time: 120 ms in all spectra). Backbone resonance assignment was done semi-automatically using CARA v1.8.4.2 and Autolink II v0.8.7 (Masse and Keller, 2005). The sidechain resonances were assigned manually. NOESY cross-peaks were picked and quantified using ATNOS (Herrmann et al., 2002)(implemented in UNIO’10 v2.0.2 (Serrano et al., 2012)). TALOS-N was used to calculate φ- und ψ-angles based on the backbone chemical shifts (Shen and Bax, 2013). The resonance assignment of paxillin LIM2/3 has been deposited to the BMRB (Entry 51154).

### Structure calculation

Initial Structure calculation was done using Cyana v3.0 (Guntert, 2004), with the protein sequence, the resonance assignment (CARA), NOESY-peaklists (ATNOS) and backbone angular restraints (TALOS-N) as input. In later stages of the calculation additional distance and angular constraints for a tetrahedral Zinc-coordination of the respective amino acids were implemented. The coordination mode of the four histidines (H403, H406, H462 and H492) was determined by the difference of chemical shifts of C^δ2^ and C^ε1^ and in all cases δ (C^ε1^) – δ (C^δ2^) was larger than 17 ppm indicating a coordination via N^δ1^ for all histidines. The structures were visualized and analyzed with PyMOL v1.3 (The PyMOL Molecular Graphics System, Version 2.0 Schrödinger, LLC). The coordinates of the final ensemble have been deposited to the PDB (Accession code: 7QB0)

**Table 1:**
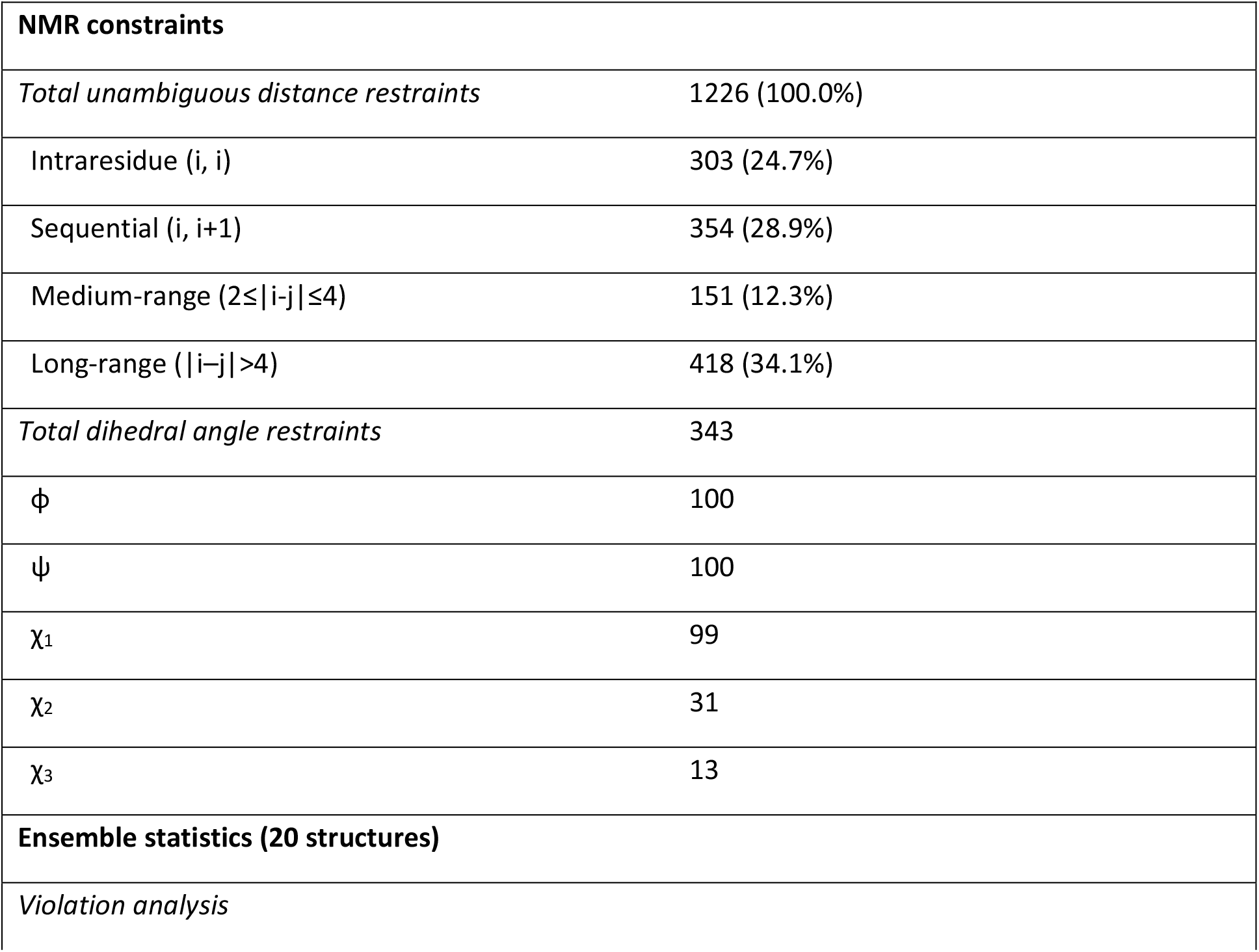
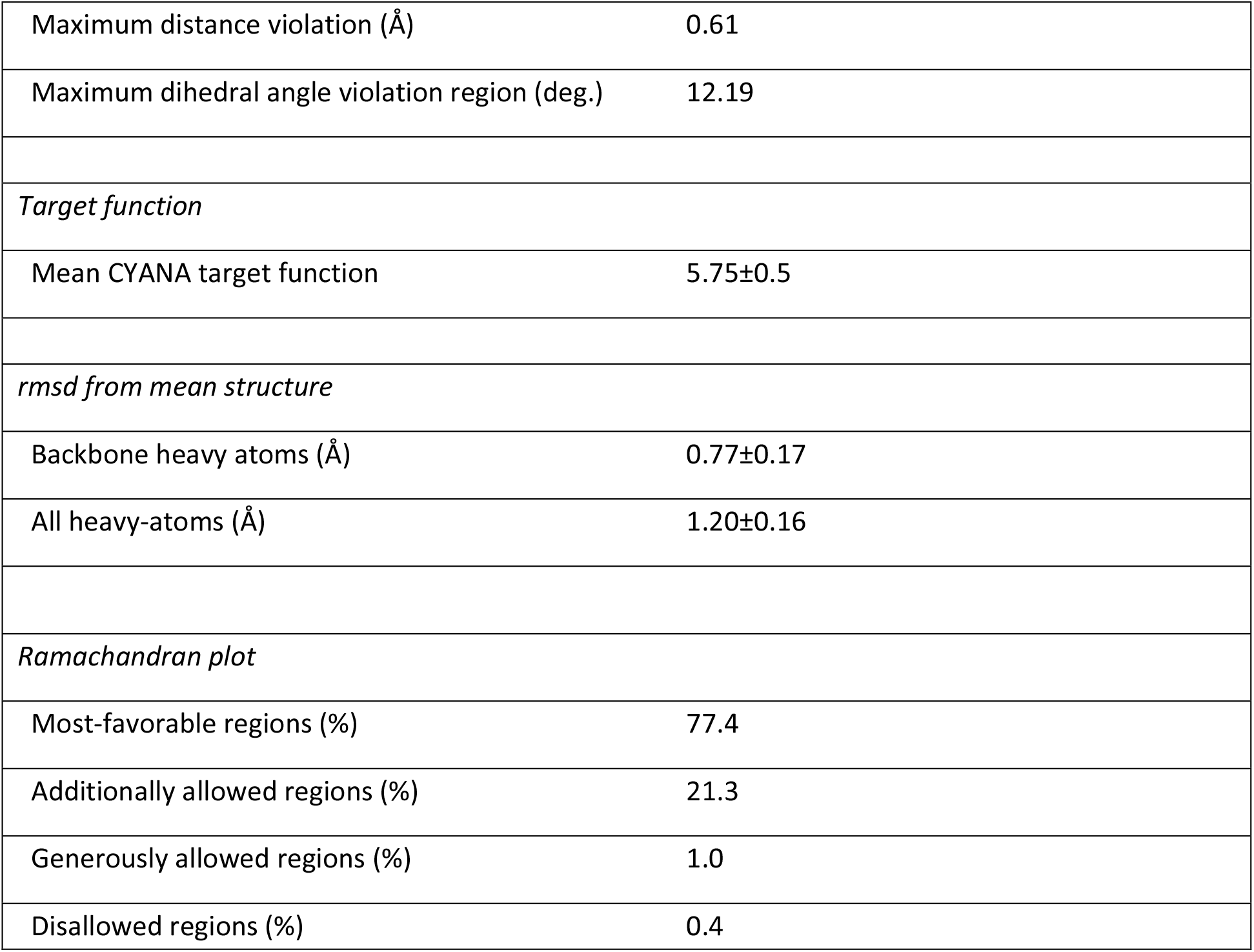
Structural statistics

|  |  |
| --- | --- |
| **NMR constraints** |  |
| *Total unambiguous distance restraints* | 1226 (100.0%) |
| Intraresidue (i, i) | 303 (24.7%) |
| Sequential (i, i+1) | 354 (28.9%) |
| Medium-range ($2 \leq \vert i-j\vert \leq 4$) | 151 (12.3%) |
| Long-range ($\vert i-j\vert > 4$) | 418 (34.1%) |
| *Total dihedral angle restraints* | 343 |
| $\phi$ | 100 |
| $\psi$ | 100 |
| $\chi_1$ | 99 |
| $\chi_2$ | 31 |
| $\chi_3$ | 13 |
| **Ensemble statistics (20 structures)** |  |
| *Violation analysis* |  |

|  |  |  |
| --- | --- | --- |
| 600 | Maximum distance violation (Å) | 0.61 |
| 601 | Maximum dihedral angle violation region (deg.) | 12.19 |
| 602 |  |  |
| 603 | *Target function* |  |
| 604 | Mean CYANA target function | 5.75±0.5 |
| 605 |  |  |
| 606 | *rmsd from mean structure* |  |
| 607 | Backbone heavy atoms (Å) | 0.77±0.17 |
| 608 | All heavy-atoms (Å) | 1.20±0.16 |
| 609 |  |  |
| 610 | *Ramachandran plot* |  |
| 611 | Most-favorable regions (%) | 77.4 |
| 612 | Additionally allowed regions (%) | 21.3 |
| 613 | Generously allowed regions (%) | 1.0 |
| 614 | Disallowed regions (%) | 0.4 |

### Chemical shift perturbation mapping

^1^H-^15^N HSQC spectra were recorded on a Bruker Avance III 600 MHz spectrometer equipped with a 5 mm BBI probe and a Bruker Avance NEO 500 MHz equipped with a H/C/N TCI CryoProbe Prodigy (Bruker Biospin GmbH, Rheinstetten, Germany). Chemical shifts were referenced to internal sodium 3-(Trimethylsilyl)propane-1-sulfonat-d6 (DSS) at 0.0 ppm. The spectra were processed and analyzed with Topspin® v2.1-4.0 (Bruker Biospin GmbH, Rheinstetten, Germany) and CARA (v. 1.9.1.5.). For NMR-experiments the proteins were concentrated by repeated ultrafiltration (Amicon Ultra-4 Ultracel-3 kDa centrifugal filter device, Merck Millipore, Burlington, USA).

Experiments were performed at 298 K in buffer containing 50 mM Na_2_HPO_4_, 150 mM NaCl, 0.1 mM ZnSO_4_, 1 mM DTT, 5% D_2_O, pH = 6.2 (if integrin was ^15^N-labeled) or 7.5 (if paxillin was ^15^N-labeled). Experimental procedure: To a sample of ^15^N-labeled protein a stock solution of unlabeled interaction partner up to a final concentration of the respective constructs was added for collecting ^1^H-^15^N HSQC spectra. The resonance assignment of integrin β1 was transferred from from BMRB entry 16159 (Anthis et al., 2009) and for integrin β3 from the BMRB entry 15552 (Oxley et al., 2008). Chemical shift change (Δδ) was calculated with the equation

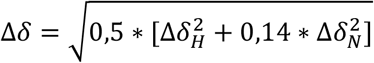

where Δδ [ppm] = δbound - δfree. The titration curves were fitted in OriginPro® (v. b9.5.5.409) using equation:

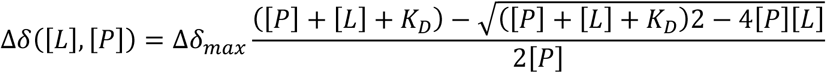

A simultaneous fit for multiple signals was used, allowing individual Δδmax values for each residue, but a global value for the dissociation constant K_D_.

### sgRNA design and cloning

For the generation of recombinant sgRNA-expression vectors, we equipped the pBluescript vector (pBS SK+, Agilent technologies, Santa Clara, CA, USA) with the murine U6 promotor controlled sgRNA expression cassette from pSpCas9(BB)-2A-GFP (PX458, a gift from Feng Zhang, Addgene plasmid # 48138) (Ran et al., 2013). Therefore, we amplified the U6 controlled sgRNA expression cassette by polymerase chain reaction (PCR) with the following primer pair: U6_sgRNA_forward: 5’-ATAGGTACCGTGAGGGCCTATTTCCC-3’ U6_sgRNA_reverse: 5’-ATACTCGAGGTCTGCAGAATTGGCGC-3’. The resulting construct was cloned into pBS SK+ via XhoI and KpnI restriction sites. The sequence verified construct (pBS-U6) was then digested with BbsI and ligated with the annealed primer pair:

MCS_oligo_forward:

5’-CACCGGGTCTTCGATGGGCCCAATTCGAATACACGTGGTTGATTTAAATGGGCCCGAAGACCT-3’

MCS_oligo_reverse:

5’-AAACAGGTCTTCGGGCCCATTTAAATCAACCACGTGTATTCGAATTGG

GCCCATCGAAGACCC-3’

to create pBS-U6 with a multiple cloning site (pBS-U6-MCS) within the BbsI restriction sites. To generate the respective pBS-U6-Cer-sgRNA plasmid, the following sgRNA Oligos Cer-KO forward: 5’-CACCGCCGTCCAGCTCGACCAGGA-3’ and Cer-KO reverse: 5’-AAACTCCTGGTCGAGCTGGACGGC-3’ were annealed and ligated into the pBS-U6-MCS vector via the BbsI restriction sites. To eliminate remaining pBS-U6-MCS after the ligation step, samples were digested with BstBI. All constructs were sequence verified by LGC Genomics.

For targeting paxillin the sgRNA oligos targeting exon 2 PXN-KO sense: 5’-CACCGACGGTGGTGGTGGGACCGG-3’ and PXN-KO reverse: 5’-AAACCCGGTCCCACCACCACCGTC-3’ were annealed and ligated into pSpCas9(BB)-2A-GFP (PX458-sgRNA mPXN)). For targeting murine integrin β3, the sgRNA oligos targeting exon 2 mITGB3-KO sense: 5’-CACCGCGGACAGGATGCGAGCGCAG-3’ and mITGB3-KO reverse: 5’-AAACCTGCGCTCGCATCCTGTCCGC-3’ were annealed and ligated into pBS-U6-MCS to generate pBS-U6-mITGB3-sgRNA.

All sgRNAs were designed with the help of the CRISPR design tool (http://crispr.mit.edu) (Hsu et al., 2013) and E-CRISP (www.e-crisp.org/E-CRISP).

### Generation of integrin β3 and paxillin knockout cell lines

For the generation of integrin β3 and paxillin knockout cell lines, Flp-In 3T3 cell line (Invitrogen) was first stably transduced with a lentiviral vector encoding Histon2B mCerulean. Therefore, human histone 2B cDNA (H2B, gift from Thomas U. Meyer, University of Konstanz, Konstanz, Germany) was amplified by polymerase chain reaction (PCR) with the following primer pair: hH2B_forward: 5’-ATAGCTAGCACCATGCCAGAGCCAGCGAAGTC-3’ hH2B_reverse: 5’-ATAACCGGTTTAGCGCTGGTGTACTTGG-3’ and cloned into pmCerulean-C1 (gift from David Piston, Vanderbilt University Medical Center, Nashville, USA) via NheI and AgeI restriction sites. The resulting construct was again subjected to PCR amplification with primers: H2B-Cer_forward: 5’-ATAGGATCCACCATGCCAGAGCCAGCGAAG-3’ and H2B-Cer_reverse: 5’-ATACTCGAGCTATTTGTACAGTTCGTCCATGCCG-3’.

The PCR product was subcloned into pWZL Blasticidin (pWZLBlast, gift from Nicole Brimer, University of Virginia, Charlottesville, USA) via BamHI and XhoI restriction sites to generate pWZLBlast-H2B-Cer. For retroviral production, 80% confluent Phoenix-Eco cells (Swift et al., 2001) were transfected with pWZLBlast-H2B-Cer and cultured for 2 days. Afterwards, the supernatant was harvested, filtered through a 0.45 µm pore-size filter unit (Minisart®, Sartorius Stedim Biotech GmbH, Göttingen, Germany) and applied on previously seeded NIH3T3 Flp-In cells at a ratio of 1:1 (vol/vol, supernatant : NIH3T3 growth medium) together with 4 µg/ml Polybrene® (Sigma-Aldrich). Transduced cells were cultured in regular growth medium supplemented with 5 µg/ml blasticidin (Carl Roth GmbH + Co. KG, Karlsruhe, Germany). Cerulean-positive cells were sorted by FACS and seeded as single cells into 96-well plates to generate clonal Cerulean-positive NIH3T3 H2B-Cer Flp-In cell lines.

Paxillin knockout cells were generated by transiently transfecting NIH3T3 H2B-Cer Flp-In cells with a combination of PX458-sgRNA mPXN + pBS-U6-Cer-sgRNA at a ratio of 1:5. Integrin β3 knockout cells were generated by transiently transfecting NIH3T3 H2B-Cer Flp-In cells with a combination of PX458-sgRNA Cer + pBS-U6-mITGB3-sgRNA at a ratio of 1:5. 10 days after transfection, single cerulean negative cells were sorted into 96 well plates and clonal cell lines were expanded and knockout of the target protein was verified by Western Blot.

### Stable complementation of knockout cells

For complementation of integrin β3, cDNA of murine integrin β3 (gift from Michael Davidson, addgene plasmid # 54130) was amplified by PCR using the following primers mITGB3 forward: 5’-GATGACACTAGTGACCGCCATGCGAGCGCAGTG-3’ and mITGB3-fl reverse: 5’-TCGGCAGCCCTCGAGCTAAGTCCCCCGGTAGGTGATATTG-3’; mITGB3 forward and mITGB3∆8aa reverse: 5’-TCGGCAGCCCTCGAGCTAGAAGGTGGAGGTGGCCTCTTTATAC-3’; mITGB3 forward and mITGB3∆3aa reverse: 5’-TCGGCAGCCCTCGAGCTAGTAGGTGATATTGGTGAAGGTGGAGGTG-3’. The respective products were cloned into pEF5/FRT-DEST (gift from Rajat Rohatgi, Addgene plasmid # 41008) using SpeI and PspXI restriction sites.

For complementation of Flp-In paxillin KO cells, we equipped the expression vector pEF5/FRT-DEST with a GFP-tag adjacent to a LoxP site for C-terminal protein tagging via Cre-Lox recombination. Therefore, the respective sequence was amplified by PCR from pEGFP C1 (Clontech, Takara Bio Europe, Saint-Germain-en-Laye, France) with the following primer pair: EGFP_forward: 5’-GCCTAGACTAGTTAGCGCTACCGGTCGCCACCATG-3’ EGFP_reverse: 5’-GCAGCGCTCGAGGGCTGATTATGATCAGTTATCTAGATCC-3’. The resulting construct was cloned into pEF5/FRT-DEST via SpeI and PspXI restriction sites to generate the expression vector pEF5/FRT EGFP C1 loxp.

The coding sequences (CDS) of paxillin was amplified by PCR with the following primers:

PXN-fl forward: 5’-ACTCCTCCCCCGCCATGGACGACCTCGACGCCCTGCTG-3’

PXN-fl reverse: 5’- CCCCACTAACCCGCTAGCAGAAGAGCTTGAGGAAGCAGTTCTGACAGTAAG

G-3’. PXN ∆LIM4 forward: 5’- ACTCCTCCCCCGCCATGGACGACCTCGACGCCCTGCTG -3’, PXN ∆LIM4 reverse: 5’-CCCCACTAACCCGCGAGCCGCGCCGCTCGTGGTAGTGC-3’; The paxillin 4A mutant was generated by overlap extension PCR. In a first PCR two fragments were generated using primers PXN-fl forward and PXN-4A reverse: 5’-TGCTGCTGCAGCGAATGGCGTGAAGCATTCCCGGCACACAAAG -3’. For the second fragment primers PXN-4A forward: 5’-GCTGCAGCAGCATTCTTCGAGCACGACGGGCAGCCCTAC -3’ and PXN-fl reverse were used. In a second step the two fragments were annealed by overlap extension PCR and amplified using primers PXN-fl forward and PXN-fl reverse.

The respective products were cloned into the pDNR-Dual-LIC vector according to the ligation independent cloning (LIC) strategy. The sequence verified constructs were then subcloned into the expression vector pEF5/FRT EGFP-C1 by Cre-Lox recombination.

Respective knockout cell lines were complemented by transient transfection of 0.8 µg cDNA coding for the gene of interest + 3.2 µg Flp recombinase expression vector (pOG44) using jetPRIME® transfection reagent (Polyplus transfection, Illkirch, France). After 3 days, positive cells were selected by addition of 250 µg/ml Hygromycin B for 8 days.

### Flow cytometry

Cells were trypsinized and suspended in growth medium. Samples were centrifuged at 100g for 3 min and the resulting pellet was re-suspended in FACS buffer (PBS with 5% FCS, 2 mM EDTA). Cells were washed once in FACS buffer and 1×106 cells per sample were incubated with monoclonal anti-integrin antibodies as indicated for 1h at 4°C under constant rotation. Cells were washed three times with FACS buffer, followed by incubation for 30 min with a Rhodamine-Red conjugated secondary antibody. Cells were analysed by flow cytometry (BD LSRFortessa, FACSDiva™ software, BD Biosciences, Heidelberg, Germany).

### Cell spreading analysis

Sterile glass coverslips were coated over night at 4°C with 5µg/ml vitronectin or 5 µg/ml fibronectin type III repeats 9-11 (FNIII9-11). Cells were starved overnight in starvation medium (DMEM + 0.5% FCS). After 12h starvation cells were trypsinized, trypsin was inactivated using soybean trypsin inhibitor (Applichem; 0.25 mg/ml in DMEM + 0.25% BSA). Cells were pelleted by centrifugation (100g, 3 min, RT) and suspended in DMEM + 0.25% BSA. Cells were kept in suspension for 30 min before seeding on coated glass coverslips. After 30 and 120 min of adherence, cells were washed once with PBS++ (0.5 mM MgCl_2_, 0.9 mM CaCl_2_), fixed with 4% PFA in PBS for 15 min at RT, washed thrice in PBS, permeabilized with 0.4% Triton-X-100 in PBS for 5 min at RT, washed thrice in PBS and blocked for 30 min in blocking buffer (10% heat inactivated CS in PBS). Cells were stained with CellMaskTM Orange (diluted to 5µg/ml in blocking buffer) and DAPI (diluted to 0.2 µg/ml in blocking buffer) for 30 min at RT. Images were analysed by a custom build ImageJ macro (Bioimaging Center, University of Konstanz)

### Fluorescent Microscopy and microscope settings

For confocal laser scanning microscopy all images were taken from fixed specimens embedded in Dako fluorescent mounting medium (Dako Inc, Carpinteria, USA) on a LEICA SP5 confocal microscope equipped with a 63.0x/1.40 NA oil HCX PL APO CS UV objective and analyzed using LAS AF Lite software. All images were acquired in xyz mode with 1024 × 1024 pixel format and 100 Hz scanning speed at 8 bit resolution. Fluorochromes used are Pacific Blue (excitation 405 nm, emission bandwidth: 435 – 475 nm); CF405M (excitation 405 nm, emission bandwidth: 435 – 475) GFP (excitation 488 nm, emission bandwidth: 500 – 525 nm); CellMask Orange (excitation 561 nm, emission bandwidth: 571 – 613 nm), RFP (excitation 561 nm. Emission bandwidth 571 – 613 nm) and Cy5 (excitation 633 nm, emission bandwidth: 640 -700 nm). Images were processed using ImageJ by applying the same brightness/contrast adjustments to all images within one experimental group.

### TIRF microscopy

Cells were starved overnight in starvation medium (DMEM + 0.5% FCS). After 12h starvation cells were trypsinized, trypsin was inactivated using soybean trypsin inhibitor (Applichem; 0.25 mg/ml in DMEM + 0.25% BSA). Cells were pelleted by centrifugation (100g, 3 min, RT) and suspended in DMEM + 0.25% BSA. Cells were kept in suspension for 30 min before seeding on Wilco dishes, coated with 5 µg/ml vitronectin. Cells were imaged with a GE DeltaVision OMX Blazev4, equipped with a 60x/1.49 UIS2 APON TIRFM objective in Ring TIRF mode. Settings were adjusted to reach clean TIRF illumination without epifluorescence. A separate sCMOS camera was used for each channel and images were later aligned using OMX image alignment calibration and softWoRx 7.0. Fluorophores used were GFP (excitation wavelength 488 nm, emission bandwidth: 528/48 nm) and Cy5 (excitation wavelength 647 nm, emission bandwidth: 683/40 nm).

### Opa-protein triggered integrin clustering (OPTIC)

OPTIC was performed as described previously (Baade et al., 2019). Briefly, 293T cells were transfected with pcDNA3.1 CEACAM3-ITGB fusion constructs together with cDNA coding for the protein of interest fused to eGFP. 48 h post-transfection, cells were seeded on coverslips coated with 10 µg/ml poly-L-lysine in suspension medium (DMEM + 0.25% BSA). After 2h, adherent cells were infected with Pacific Blue-stained *Neisseria gonorrhoeae* (Opa52-expressing, non-piliated *N. gonorrhoeae* MS11-B2.1, kindly provided by T. Meyer, Berlin, Germany) at MOI 20 for 1h in suspension medium. After 1h cells were fixed for 15 min with 4% paraformaldehyde in PBS at room temperature followed by 5 min permeabilization with 0.1% Triton X-100 in PBS. After washing with PBS, cells were incubated for 10 min in blocking solution (10% heat inactivated calf serum in PBS) and stained for CEACAM3. After washing, cells were again incubated for 10 min in blocking solution followed by secondary antibody staining. Coverslips were mounted on glass slides using Dako fluorescent mounting medium (Dako Inc, Carpinteria, USA).

## Acknowledgments

This work was funded by Deutsche Forschungsgemeinschaft via CRC969, project B06 (to CRH) and via RTG 2473 “Bioactive Peptides”, project C3 (to HMM). We gratefully acknowledge initial contributions to peptide and protein preparation and interaction studies by J. Ude, M. Roth, M. Gallandi, and S. Feindler-Boeckh, as well as expert support in mass spectrometry by Dr. I. Starke. We thank D. Schlaepfer (UCSD, San Diego, CA) for providing GFP-FAK re-expressing FAK-/- murine fibroblasts and R. Fässler and R. Böttcher (MPI for Biochemistry, Martinsried, Germany) for providing kindlin1/kindlin2 deficient mouse fibroblasts. We would also like to thank the Core Facilities of the University of Konstanz for excellent help and support with cell sorting (A. Sommershof, FlowKon) and microscopy (M. Stöckl, Bioimaging Center).

## Author contributions

H.M.M. and C.R.H. conceived the study. H.M.M., C.R.H., T.B., M.M., A.P., C.P. and N.K. designed the experiments. T.B., C.P. and L.S. cloned constructs, established cell lines and performed OPTIC assays. T.B. performed cell-spreading experiments, TIRF microscopy and pulldown assays. M.M., A.P., N.K. and R.N. cloned and prepared paxillin and integrin proteins and peptides and performed NMR experiments. All authors analyzed and interpreted data. T.B., M.M., H.M.M. and C.R.H wrote the manuscript.

