## Supplementary Figures for "A flexible loop in the paxillin LIM3 domain mediates direct binding to integrin β3"

#### Supplementary Figure legends

**Figure S1: Paxillin and closely related LIM-domain proteins localize to clustered integrin-β1 or -β3 ct.**

(A) Schematic overview of the OPTIC workflow. Opa-expressing Ngo are used to cluster CEACAM3-integrin β cytoplasmic tail fusion proteins potentially resulting in the recruitment of an intracellular protein of interest (POI). (B) 293T cells were transiently co-transfected with a CEACAM3-ITGB1 (ITGB1) or CEACAM3-ITGB3 (ITGB3) fusion construct together with GFP labelled cytosolic proteins and seeded on poly-L-lysine. Cells were infected for 1 h with Pacific Blue labelled Neisseria gonorrhoeae (Ngo), fixed, and stained for CEACAM3. Bars represent 2 µm. (C) Quantification of A. Each data point reflects the recruitment ratio R in a CEACAM3-expressing cell with associated bacteria. Horizontal lines indicate mean values and 95% confidence intervals (whiskers) of n = 60 cells from three independent experiments. Statistical significance was calculated using one-way ANOVA, followed by Bonferroni Multiple Comparison Test (*** $p<0.001$, ns = not significant). (D) Western blots of whole cell lysates of transfected HEK cells from the experiment in Suppl. Fig. 1B. Membranes were probed with a monoclonal antibody against GFP. Coomassie staining was used to verify equal loading of the membrane.

**Figure S2: Zinc fingers of the LIM3 domain show increased structural flexibility and are highly conserved across species**

(A) Heteronuclear $^{15}$N[$^{1}$H] NOE. The intensity ratio between spectra with and without $^{1}$H saturation is displayed vs residue number. Values of 0.8 indicate rigid parts of the structure. Values smaller than 0.8 indicate increasing flexibility on the ps-to-ns timescale. (B) Sequence alignment of paxillin LIM3 domain across different species: Alignment was performed using the structural alignment tool from T-Coffee and coloured using the BoxShade tool at ExPASy. Identical residues are shaded in black; highly similar residues are shaded gray.

**Figure S3: Distinct sequence motifs within integrin-β1 and -β3 are crucial for recognition by paxillin.**

(A) $^{15}$N-HSQC titration of 300 μM $^{15}$N integrin β1 ct (ITGB1 ct) with paxillin LIM2/3 (PXN LIM2/3). Paxillin was added in concentrations up to 500 μM. Boxes show a selection of signals affected by CSPs (residues K784, T788 & T789) in the presence of 0 μM (black), 150 μM (green), 300 μM (blue) and 650 μM (red) paxillin LIM2/3. Insets show the concentration dependence of combined amide CSPs globally fitted to a one site binding model. (B) Combined amide CSPs of 300 μM $^{15}$N integrin β1 ct in the presence of 650 μM paxillin LIM2/3 vs residue number of integrin β1 ct. (C) $^{15}$N-HSQC titration of 300 μM $^{15}$N paxillin LIM2/3 (PXN LIM2/3) with integrin β3 ct Δ8aa (ITGB3 Δ8aa). Integrin was added up to a concentration of 600 μM. (D) Combined amide CSPs of 300 μM $^{15}$N paxillin LIM2/3 in the presence of 600 μM integrin β3 ct Δ8aa vs residue number of paxillin LIM2/3. (E) 293T cells were transiently co-transfected with a CEACAM3-3 ITGB3 (CEA3-ITGB3) fusion construct or truncated mutants together with GFP labelled paxillin wt and seeded on poly-L-lysine. Cells were infected for 1 h with Pacific Blue-labelled Neisseria gonorrhoeae (Ngo), fixed, and stained for CEACAM3. Bars represent 1 μm. (F) Quantification of paxillin recruitment to CEA3-ITGB3 clusters. Shown are means and 95% confidence intervals of n=60 cells from three independent experiments. Significance was calculated using one-way

ANOVA followed by Bonferroni Multiple Comparison Test. Significance levels compared to paxillin wt are indicated (ns: not significant; *** p≤0.0001; ** p≤0.01).

**Figure S4: Complementation of integrin β3 deficiency of Kindlin KO cells**

(A) Flow cytometric analysis of integrin β3 knockout and re-expression cell lines. Cells were stained with a monoclonal antibody against mouse integrin β3. (B) WCL of integrin β3 cell lines. Equal number of cells were lysed. Tubulin was used as loading control. (C) Flow cytometric analysis of Kindlin 1/2 knockout ($Kind_{KO}$) and Kindlin 1/2 flox cells ($Kind_{Ctrl}$) stained for different integrins. (D) $Kind_{KO}$ cells were transduced with either murine full-length integrin β3, truncated integrin β3 Δ8aa or Δ3aa or empty vector backbone (mock). Cells were analysed for their surface expression of various integrin subunits using flow cytometry. (E) Serum starved $Kind_{KO}$ cells stably expressing full length integrin β3 or truncated mutants were seeded on glass coverslips coated with 50 µg/ml vitronectin or poly-Lysin for 4 h. Cells were fixed and stained for endogenous talin.

**Figure S5: Titration of $^{15}$N-labeled paxillin LIM2/3 with integrin-β3 ct.**

To validate the dissociation constant $K_D$ determined by the inverse titration shown in Fig. 2, $^{15}$N paxillin was titrated with unlabeled integrin-β3 ct. Titration curves were obtained by globally fitting the data to a one site binding model, and selected curves are displayed.

**Figure S6: NMR experiments with paxillin point mutants show that some residues in the flexible loop are essential for maintaining a stably folded structure.**

Alanine scan of paxillin LIM3's flexible loop region. Shown are superpositions of $^{1}H$-$^{15}N$-HSQC spectra of wildtype paxillin LIM2/3 (black) and paxillin LIM2/3 mutants (red). (A) Paxillin LIM2/3 F475A. (B) Paxillin LIM2/3 V476A. (C) Paxillin LIM2/3 G478A. (D) Paxillin LIM2/3 S479A. (E) Paxillin LIM2/3 F480A. (F) Paxillin LIM2/3 F481A.

**Figure S7: The loop mutant paxillin LIM2/3 4A does no longer bind to integrin β3 ct.**

(A) $^{15}N$-HSQC titration of 300 µM $^{15}N$ paxillin LIM2/3 4A (PXN LIM2/3 4A) with integrin β3. Integrin was added in concentrations up to 1582 µM. Boxes show a selection of signals affected by CSPs (residues F421 & Y453) in the presence of 0 µM (black), 632 µM (green), 1107 µM (blue) and 1582 µM (red) PXN LIM2/3 4A. (B) Combined amide CSPs of 250 µM $^{15}N$ PXN LIM2/3 4A in the presence of 630 µM integrin β3 vs residue number of paxillin. Lines indicate average $\Delta\delta$+ 1x s.d. (yellow), $\Delta\delta$ + 2x s.d. (orange) and $\Delta\delta$ + 3x s.d. (red).

### Supplementary Figures

Baade et al. Supplementary Figure 1

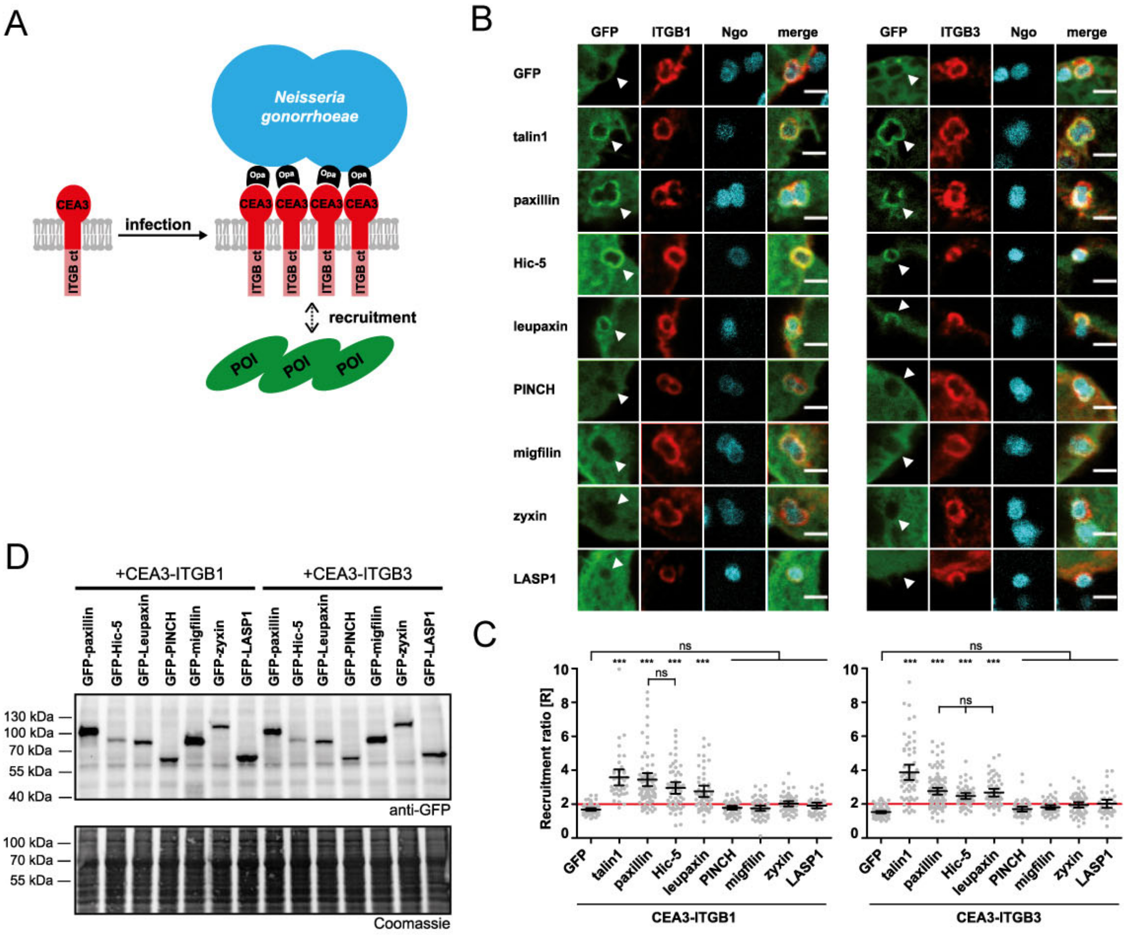

**Baade et al. Supplementary Figure 2**

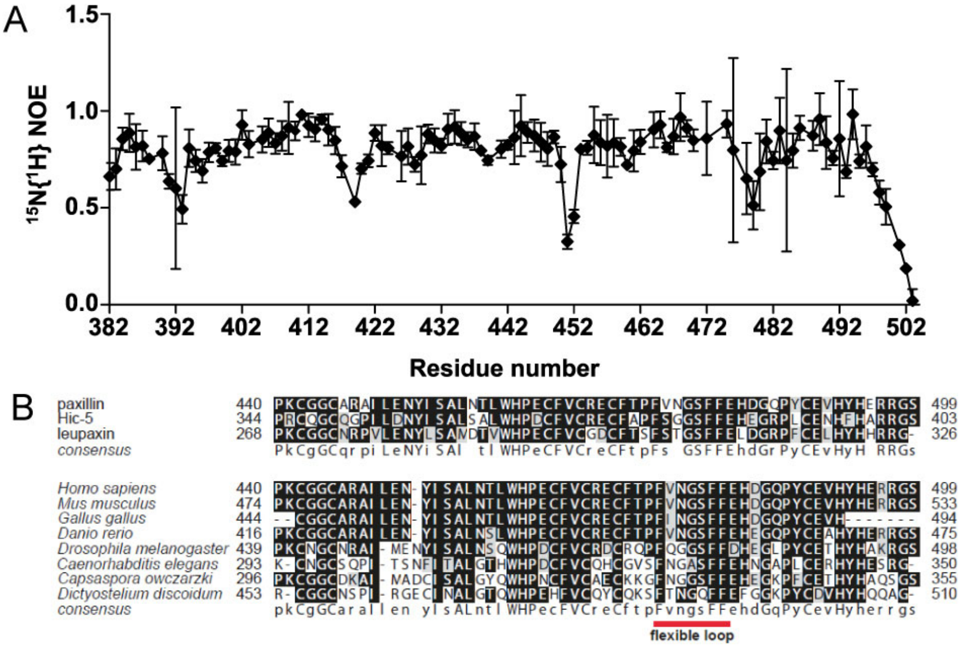

#### Baade et al. Supplementary Figure 3

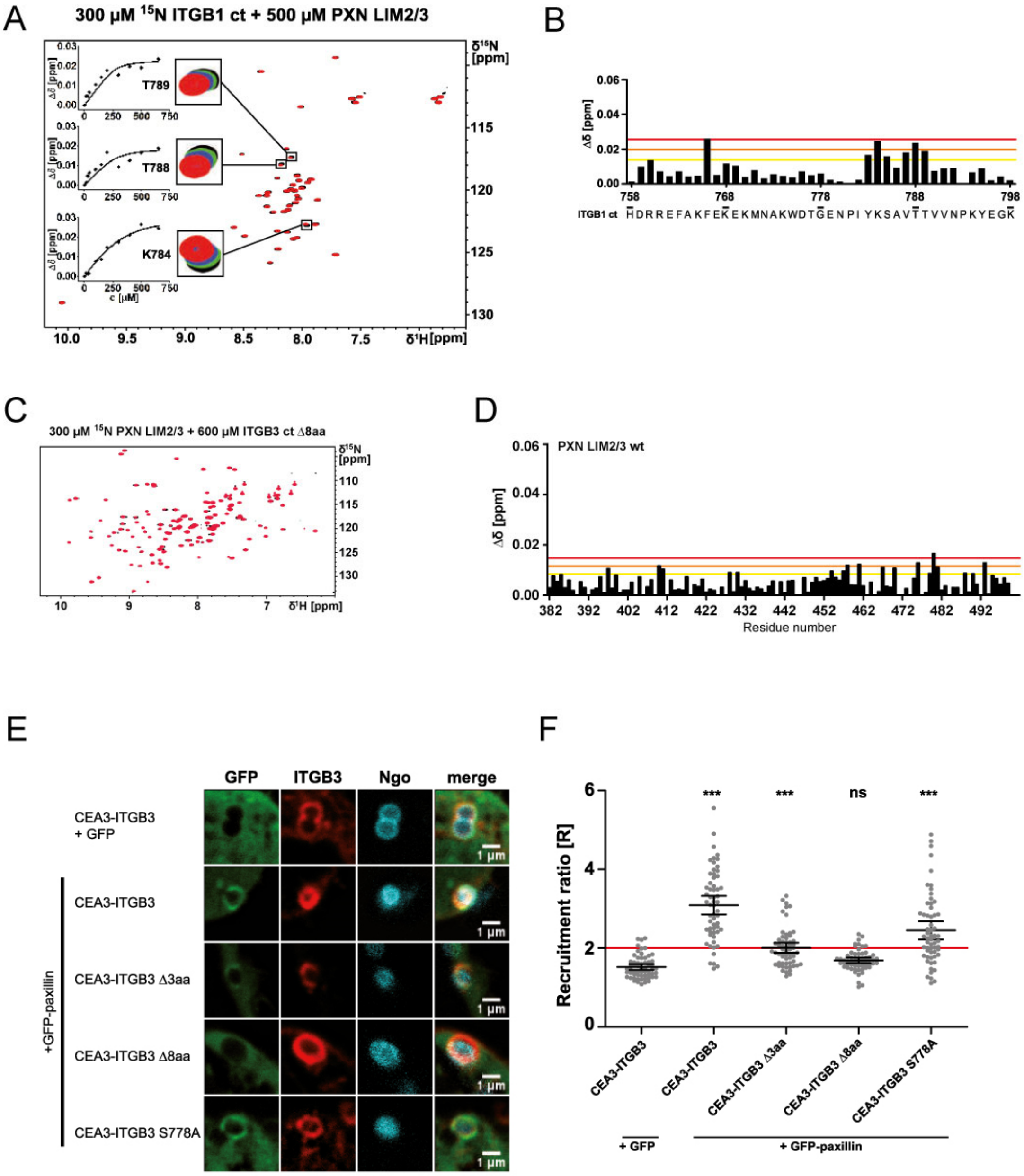

#### Baade et al. Supplementary Figure 4

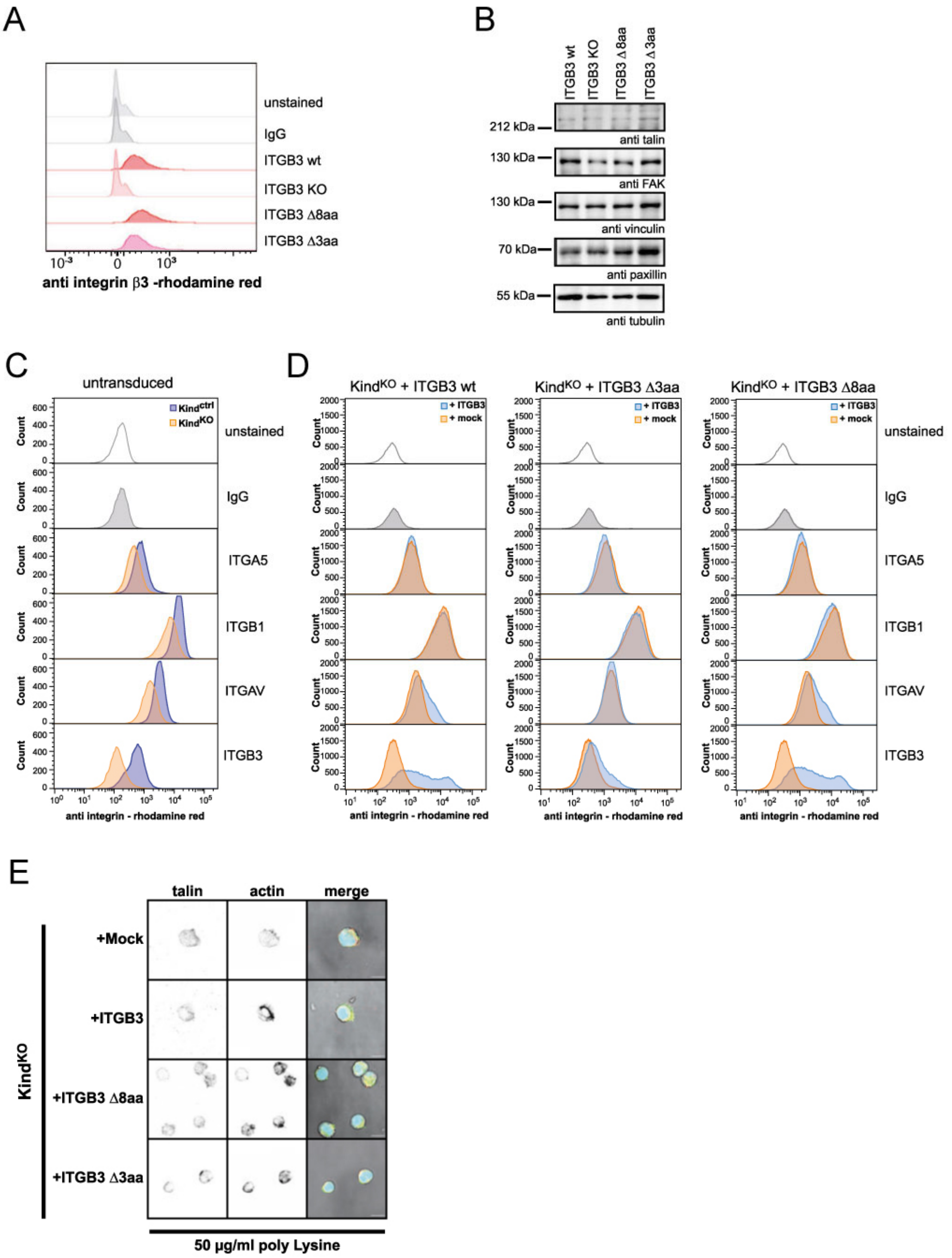

Baade et al. Supplementary Figure 5

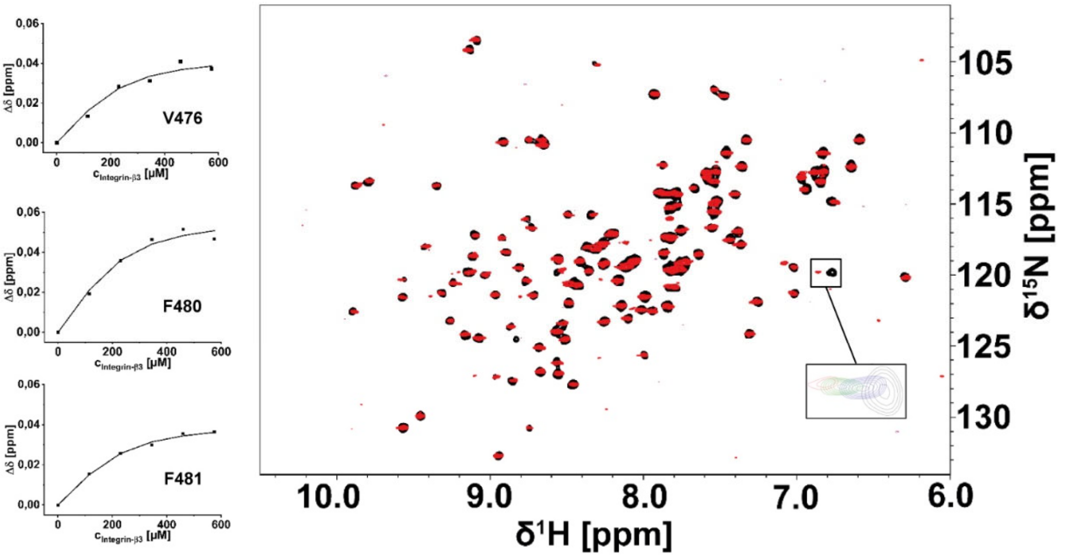

**Baade et al. Supplementary Figure S6**

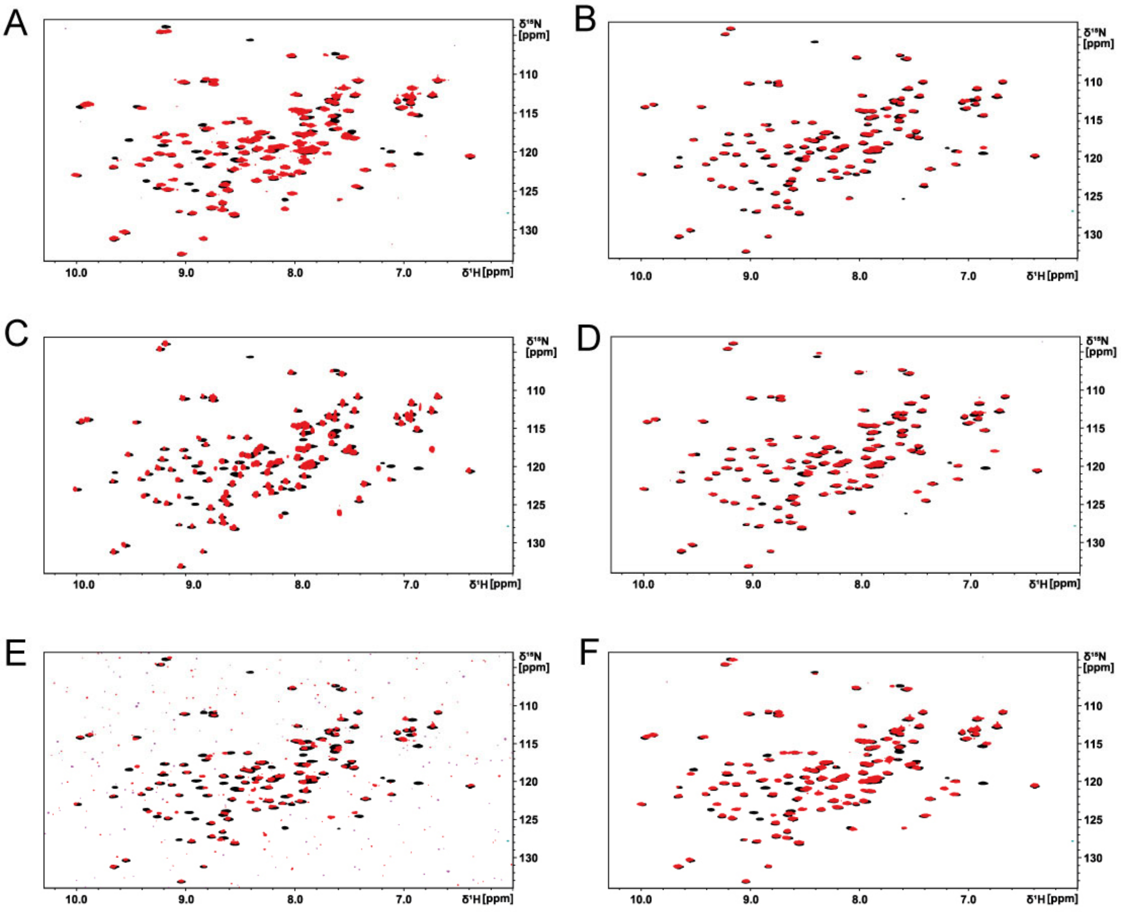

**Baade et al. Supplementary Figure 7**

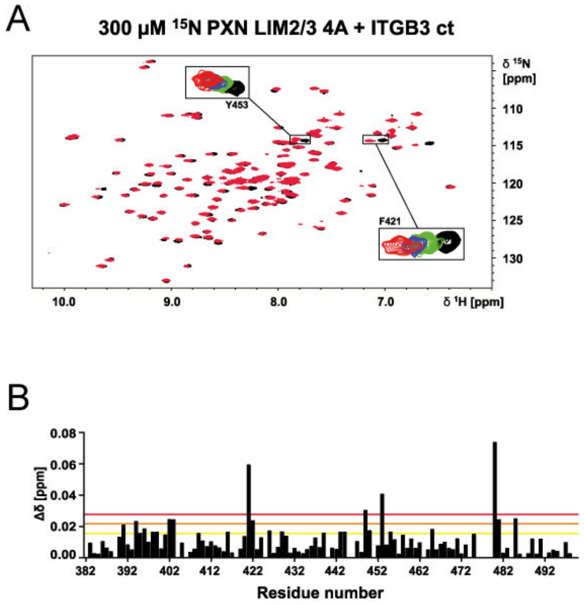
